# Distinct transcriptional responses to mild cold versus warm temperatures in adult *Drosophila melanogaster* ovaries

**DOI:** 10.64898/2026.09.24.754180

**Authors:** Ana Caroline P. Gandara, Zita Y. Gao, Yuh Chwen G. Lee, Daniela Drummond-Barbosa

## Abstract

Temperature influences fertility across diverse organisms, yet the mechanisms underlying how suboptimal temperatures affect gamete production and quality remain largely unknown. We previously showed that chronic exposure of adult *Drosophila melanogaster* females to mild cold promotes the maintenance of germline stem cells (GSCs) and high oocyte quality over time despite reducing the rates of oogenesis, while exposure to warm temperature causes death of early germline cysts and vitellogenic follicles and a severe decrease in oocyte quality. To explore potential mechanisms underlying these highly distinct responses, we compared the ovarian transcriptomes of females maintained at these temperatures (18°C or 29°C) to that of 25°C controls. We found that 18°C upregulates or downregulates ∼2.5 times as many genes as 29°C, indicating that the ovary mounts active physiological responses to mild cold and warm temperatures—as opposed to simply undergoing passive changes driven by thermodynamics.

Gene set enrichment analysis revealed modulation of genes involved in neuronal signaling in opposite directions at 18°C versus 29°C. Most genes, however, exhibit temperature-specific regulation: 29°C upregulates synaptic transmission genes and downregulates lipid biosynthesis genes, whereas 18°C upregulates actin cytoskeleton genes and downregulates cell adhesion and lipid organization genes. Notably, mild cold or warm temperature specifically modulated (either up or down) the expression of distinct sets of transposable elements (TEs), suggesting the existence of temperature-dependent TE regulatory mechanisms and/or downstream effects. Finally, we show that GSCs at 18°C have increased retrotransposon *R2* transcript levels, larger nucleolar size, and elevated levels of the known stemness factor phosphorylated Mad, leading to a working model whereby elevated ribosome biogenesis supports increased stemness signaling to promote GSC maintenance in mild cold. These findings suggest potential mechanisms and open new questions for investigation towards a deeper understanding of how temperature modulates gene expression and impacts germline development and quality—which are essential for the perpetuation of species.

## Introduction

Organisms are constantly exposed to changes in environmental factors, including temperature. Cold-blooded (poikilothermic) animals are particularly susceptible to changes in environmental temperature. Indeed, over the past several decades, global warming, combined with intensive agricultural land use and loss of habitat, has led to major reductions in insect overall abundance and number of species (1–3). In particular, the effects of climate change have been overwhelmingly negative for insect reproduction, causing changes in population size (4).

Although warm-blooded (homeothermic) animals have elaborate thermoregulation mechanisms, their body temperature varies with body site, age, circadian rhythm, and hormonal status (5) and can be disrupted by extreme environmental temperatures, fever, or neurologic disorders (6).

Notably, heat stress impairs ovarian function in humans and other mammalian species (7,8). For example, heat stress disrupts metabolism and oocyte growth, maturation, and developmental competence in livestock (9,10). Therefore, elucidating the mechanisms underpinning the effects of temperature on reproduction is broadly relevant.

*Drosophila melanogaster* is a well-established model for investigating the regulation of oogenesis by environmental factors (11). Adult *D. melanogaster* females have a pair of ovaries with 15–20 ovarioles each, and each ovariole has an anterior germarium housing two to three germline stem cells (GSCs) within a niche formed by cap cells (Fig 1A-1C). The niche produces bone morphogenetic protein (BMP) signals that activate receptors on GSCs to promote stemness through phosphorylation of the transcriptional factor Mad (12). GSCs divide asymmetrically to produce a replacement GSC and a cystoblast that gives rise to a 16-cell germline cyst, which is then enveloped by follicle cells to form a follicle (also known as egg chamber). The follicle leaves the germarium and develops through 14 stages of oogenesis to produce an elongated mature oocyte (11) (Fig 1B and 1C). In contrast to previous studies focused on developmental exposure to elevated temperatures or adult exposure to high or low temperature extremes (11), we recently delineated the cellular effects of chronic exposure to suboptimal temperatures on oogenesis in adult *D. melanogaster* (13). We showed that GSC numbers and oocyte quality remain higher over time despite lower oogenesis rates in adult females maintained at 18°C (mild cold) compared to controls at 25°C, while chronic exposure to 29°C (warm) causes elevated germline death at specific stages and reduces oocyte quality (13). Temperature also influences oogenesis in many other insect species, which has broad implications given the critical roles insects play in agriculture, vector-borne diseases, and ecological balance (11).

**Fig 1.**
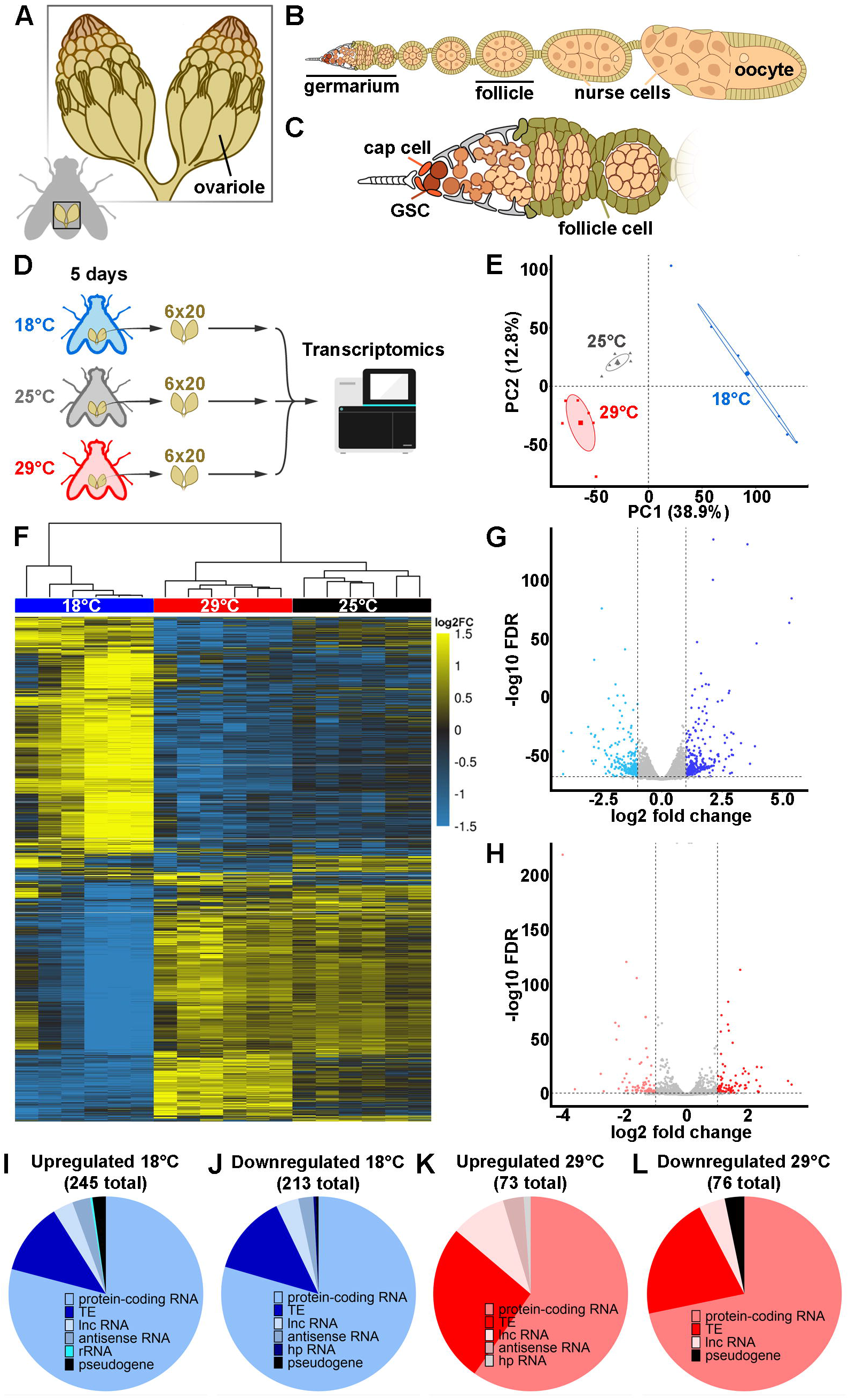
Warm and mild cold temperatures induce distinct transcriptomic responses in *D. melanogaster* ovaries. (A) Diagram of a pair of *D. melanogaster ovaries*, composed of ovarioles. (B) Diagram of an ovariole, composed of a germarium followed by developing follicles (i.e., egg chambers), each comprising a germline cyst—15 nurse cells and one oocyte— surrounded by follicle cells. (C) Diagram of germarium housing germline stem cells (GSCs; red) in direct contact with cap cells (dark orange) in the niche, mitotically dividing early GSC daughters (light orange), and 16-cell cysts (lightest orange) that become enveloped by follicle cells (green) to form new follicles. (D) Flow chart of transcriptomics sample preparation from females raised at 21-23°C and incubated (shortly after eclosing) at 18°C, 25°C or 29°C (with males) for five days prior to dissection of six replicates of 20 pairs of ovaries per temperature. (E) Principal component analysis graph showing intra-group clustering and intergroup separation and indicating gene expression correlation among different temperature conditions. Elliptical shapes indicate 95% confidence interval. (F) Heatmap of transcripts significantly (*q*≤0.05) modulated by temperature. Expression level is scaled per gene in each row, with yellow and blue representing upregulated and downregulated genes, respectively. (G and H) Volcano plots of modulated genes at 18°C (G) or 29°C (H), relative to 25°C control. Blue (G) and red (H) dots represent significantly (*q*≤0.05) modulated genes. (I-L) Pie charts showing the types of transcripts significantly upregulated (I) or downregulated (J) at 18°C, or upregulated (K) or downregulated (L) at 29°C. TE, transposable element; lnc RNA, long non-coding RNA; hp RNA, hairpin RNA; rRNA, ribosomal RNA.

Despite its importance, relatively little is known about how the ovaries of adult *D. melanogaster* or other insects respond to suboptimal temperatures at the molecular level. In *D. melanogaster*, previous studies characterized gene expression changes in the ovary during diapause (induced by incubating newly eclosed flies at 11-12°C and short photoperiod for three or four weeks) (14–16) or of whole flies in response to acute 34°C heat (17) or lower than 4°C cold stress (18–20). A few studies in other insects either during diapause or in response to chronic heat stress have also been conducted (21–23). A previous study comparing the ovarian transcriptomes of *D. melanogaster* females maintained at 29°C for two days with those of females maintained at 18°C for five days revealed temperature-dependent differences in the abundance of microRNAs and their target transcripts, as well as Piwi-interacting RNAs (piRNAs) and their cognate transposable elements (TEs) (24). However, the differences in incubation time at 29°C versus 18°C and the lack of 25°C controls make it challenging to evaluate those data.

In this study, we took advantage of the highly distinct responses of the *D. melanogaster* ovary to suboptimal temperatures to explore potential molecular mechanisms underlying these responses. We compared the ovarian transcriptome following incubation of females for five days at 18°C, 25°C, or 29°C. We found that 18°C upregulated or downregulated ∼2.5 times as many genes as 29°C, indicating a robust physiological response to mild cold. Our analysis revealed a subset of genes modulated in opposite directions at 18°C versus 29°C (e.g., neuronal signaling genes), as well as a vast majority of genes exhibiting temperature-specific regulation (e.g., lipid biosynthesis genes downregulated at 29°C, or actin cytoskeleton genes upregulated at 18°C).

Further, distinct sets of TEs were differentially expressed at 18°C versus 29°C—e.g., the *R2* non-long terminal repeat (non-LTR) retrotransposon was uniquely upregulated at 18°C— suggesting temperature-specific mechanisms of TE modulation. Finally, direct examination of GSCs showed increase in *R2* transcript numbers, nucleolar size, and phosphorylated Mad (pMad) levels specifically at 18°C relative to 25°C, suggesting that the improved maintenance of GSC at 18°C stems from enhanced nucleolar function (possibly in association with increased *R2* expression) and BMP signaling. These findings contribute new insight into how temperature controls gene expression and influences the development and quality of the germline—an immortal lineage that ensures perpetuation of all sexually reproducing species (25).

## Results and discussion

### Mild cold and warm temperatures elicit distinct ovarian transcriptomic responses in *D. melanogaster*

Suboptimal warm and cold temperatures have very distinct effects on *D. melanogaster* oogenesis (13). To identify the potential mechanisms underlying these responses, we performed RNA sequencing analysis of whole ovaries from adult females maintained for five days at 25°C (control), 18°C (mild cold), or 29°C (warm) (Fig 1D). The ovaries have indistinguishable morphology at these three temperatures at this early timepoint (13). The RNA sequencing analysis identified a total of 11,864 transcripts and 23,487 splice variants across all samples (S1 Table). To assess the effects of temperature on the ovarian transcriptome, we first performed principal component analysis (PCA). Both the transcript-level and splice variant-level transcriptomes showed intra-group clustering and significant intergroup separation, most notably at 18°C (Fig 1E and S1A Fig). Clustered heatmaps of genes that were significantly modulated by temperature indicated reproducibility among replicates and temperature-dependent modulation of many transcripts and splice variants, again mostly at 18°C (Fig 1F and S1B Fig). Accordingly, a much larger number of genes were differentially expressed at 18°C than at 29°C, compared to 25°C. Genes modulated at 18°C included 2,277 upregulated and 2,253 downregulated, with 245 and 213 modulated at least 2-fold, respectively (Fig 1G and S1 Table). At 29°C, 889 genes were significantly upregulated, and 988 genes were downregulated, with 73 and 76 genes modulated by at least two-fold, respectively (Fig 1H and S1 Table). Most differentially expressed transcripts corresponded to protein-coding genes, although transcripts for TEs, long non-coding RNAs, pseudogenes, and other RNA subtypes were also regulated by temperature (Fig 1I-1L).

### Mild cold and warm temperatures distinctly regulate heat shock response genes

The historically termed ’heat shock response’ is an evolutionarily conserved response to a variety of stressors, including temperature (26,27), and involves the activation of heat shock transcription factors (HSF), which induce the expression of heat shock proteins (HSPs)— molecular chaperones that maintain protein homeostasis (27). Accordingly, *Heat shock factor* (*Hsf*), *Heat shock protein 68* (*Hsp68*), *Hsp22*, *Hsp23*, and *Heat shock protein 70 cognate 5* (*Hsc70-5*) were significantly upregulated at 29°C and downregulated at 18°C (Fig. 2). *DnaJ-like- 1* (*DnaJ-1*) and *DnaJ-like-2* (*Droj2*), both of which encode homologs of the mammalian DNAJ/HSP40 family of proteins (cochaperones HSP70 proteins) (28), were similarly regulated (Fig 2). *D. melanogaster Hsf* mutants fail to induce *Hsp70* expression at 30°C, 33°C, or 36°C and have compromised thermotolerance (29). *Hsf* is also required in the female germline for follicle development past early pre-vitellogenic stages under control conditions (29), suggesting heat-dependent and -independent roles in oogenesis. Hsp68, which is part of the Hsp70 family of ATP-dependent chaperones (26), is induced in adult females exposed to 37°C for 60 minutes and remains stable for at least 12 hours following the heat shock (30), but its function in the ovary remains unknown. The ATP-independent small heat shock proteins Hsp22 and Hsp23 are expressed in distinct somatic cell types in testes in heat-shock-dependent and independent manners, respectively (31). A more recent study showed that *Hsp23* null flies recover more slowly following cold exposure (9 hours at 0°C) but are more tolerant to heat stress (18 hours at 35°C) and also appear to lay more eggs than wildtype controls at 27°C, but not 19°C or 25°C (32), suggesting complex roles for Hsp23. Overexpressed *D. melanogaster* Hsp22 was shown to physically interact with Hsp70 and several ATP synthase subunits and increase oxygen consumption and ATP production in cultured human cells (33). Moreover, ubiquitous overexpression of *Hsp22* in flies causes increased lifespan and upregulation of mitochondrial *Hsp60*, *Hsp70*, and proteins involved in the tricarboxylic acid cycle and electron transport chain, suggesting a role in mitochondrial stress response (34). Interestingly, *Hsp60A*, *Hsp67Ba*, *Hsp70Ab*, and *DnaJ homolog* (*DnaJ-H*) were specifically upregulated at 18°C (Fig 2 and S1Table), suggesting that different heat shock response genes might have specialized roles at warm versus mild cold suboptimal temperatures. Further studies will be needed to determine the potential roles of these modulated heat shock response genes (and the clients they regulate) in the ovarian response to different temperatures.

**Fig 2.**
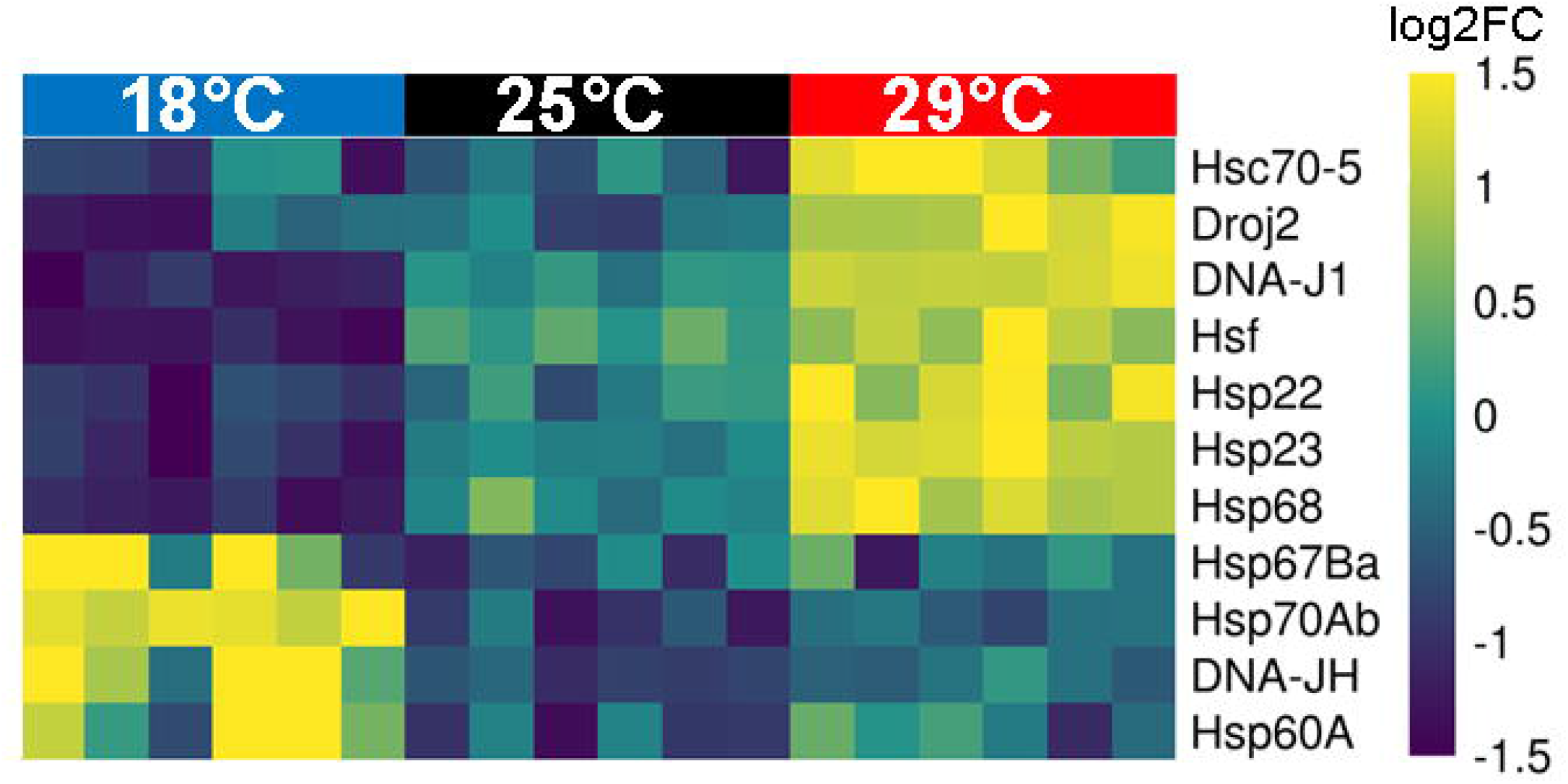
Heat shock response genes are distinctly regulated at 29°C versus 18°C. Heatmap showing heat shock response genes that are significantly (*q*≤0.05) modulated by 29°C or 18°C. Yellow and blue represent upregulated and downregulated genes, respectively.

### The *D. melanogaster* ovary actively responds to mild cold and warm temperatures through distinct transcriptomic signatures

To compare the transcriptomic patterns of the temperature responses in a systematic and unbiased manner, we first separated all 5,207 modulated genes into three categories (Fig. 3): I) regulated in opposite directions at 18°C and 29°C; II) modulated in a similar manner in both temperatures; and III) uniquely modulated at either temperature. Approximately 21% (1,098) of all modulated genes were modulated in opposite directions: 552 upregulated at 29°C and downregulated at 18°C, and 546 downregulated at 29°C and upregulated at 18°C (Fig 3A and 3B and S2 Table). Only 2% (102) of all modulated genes showed non-specific responses to temperature: 31 upregulated and 71 downregulated at both 18°C and 29°C (Fig 3C and 3D and S2 Table). Remarkably, 77% (4,006) of modulated genes were part of unique temperature-dependent responses (i.e., regulated at either 18°C or 29°C): 677 genes were modulated only at 29°C (306 upregulated and 371 downregulated) and 3,329 genes at 18°C (1,700 upregulated and 1,629 downregulated) (Fig 3 and S3 Table). When restricting our analysis to the 549 genes that are modulated at least two-fold by temperature, we found that around 9.6% (53) were regulated in opposite directions, 0.9% (5) in the same manner, and 65% (355) had unique modulation at 18°C or 29°C (Fig 3 and S2 and S3 Tables). These results support the conclusion that the ovarian responses to 18°C and 29°C are not simply the result of a global decrease or increase in gene expression driven by thermodynamic effects. Rather, the ovary actively responds to both temperatures by upregulating and downregulating distinct sets of transcripts in a highly specific manner.

**Fig 3.**
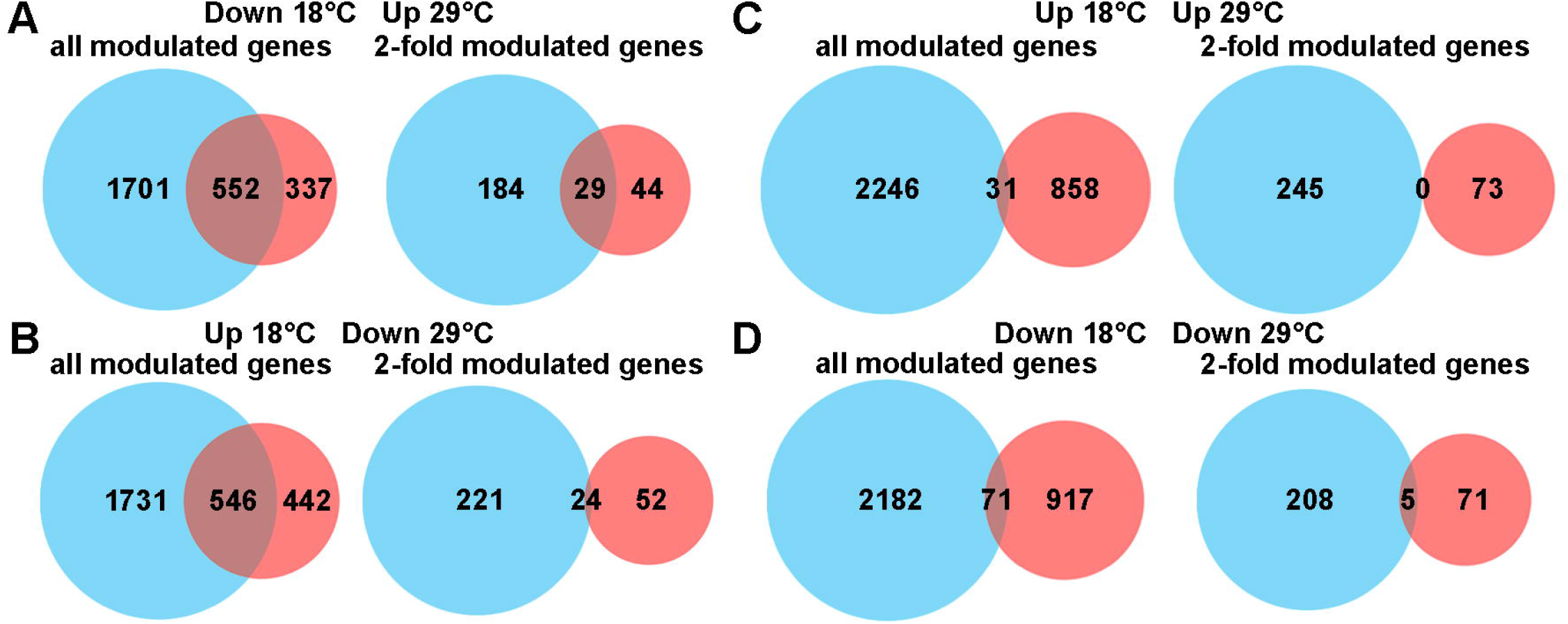
Warm and mild cold temperatures induce highly distinct ovarian transcriptomic signatures. (A-D) Venn diagrams of all modulated genes or genes modulated by at least two-fold by temperature, illustrating overlap among genes regulated in opposite directions (A and B) or in the same direction (C and D) by 18°C versus 29°C (relative to 25°C control).

### Mild cold and warm temperatures regulate neuronal and G-protein coupled receptor signaling genes in opposite directions

To identify specific biological pathways reciprocally regulated by warm and mild cold temperatures, we combined genes showing at least two-fold upregulation at 29°C and downregulation at 18°C with those displaying the reverse pattern and conducted gene set enrichment analysis (GSEA), using the recently developed bioinformatics tool PANGEA (<u>PA</u>thway, <u>N</u>etwork and <u>G</u>ene-set <u>E</u>nrichment <u>A</u>nalysis) (35) (S2 Table). The top 10 significantly enriched "GO Biological Processes" had significant overlap and fell into the broad category of neuronal and G-protein coupled receptor (GPCR) signaling (Fig 4A and S2 and S4 Tables). We used a heatmap to visualize all the reciprocally modulated genes that fell into this broad category (Fig 4B).

**Fig 4.**
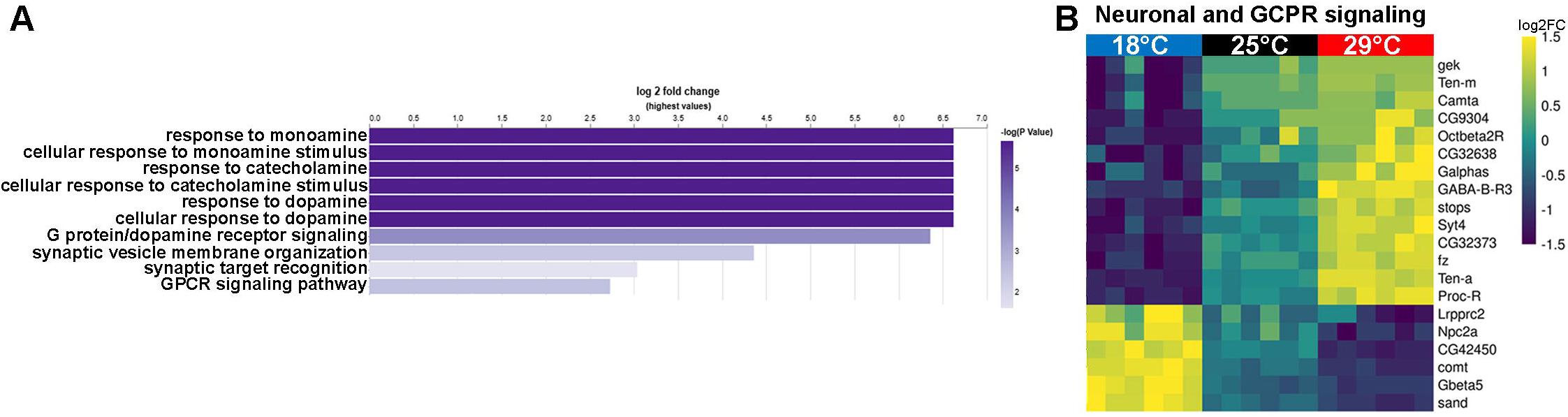
"Neuronal and G-protein coupled receptor signaling" genes are reciprocally modulated by mild cold and warm temperatures. (A) Bar graph showing the 10 most significantly (*p*≤0.05) enriched "GO Biological Processes" among all genes showing at least two-fold upregulation at 29°C and downregulation at 18°C or downregulation at 29°C and upregulation at 18°C. Altogether, these 10 most enriched "GO Biological Processes" have significant overlap (S4 Table) and fall into a broadly defined category of "neuronal and GPCR signaling". (B) Heatmap showing all reciprocally modulated genes (*q*≤0.05) involved in neuronal and GPCR signaling (S4 Table). Yellow and blue represent upregulated and downregulated genes, respectively.

Among genes upregulated at 29°C and downregulated at 18°C were *Proctolin receptor (Proc-R), Octopamine* β*2 receptor (Oct*β*2R), and metabotropic GABA-B receptor subtype 3 (GABA-B-R3)* (Fig 4B and S1 and S4 Tables), which encode GPCRs involved in neurotransmitter or neuropeptide signaling (36). Proctolin (produced from a precursor encoded by *Proc*) is an insect neuropeptide that activates Proc-R. Proctolin is degraded by Dipeptidase III (encoded by *DppIII*) to limit its activity; notably, *DppIII* was significantly upregulated at 18°C in our ovary transcriptomic dataset (S1 Table). Proctolin is primarily known as a neuromodulator that acts at neuromuscular junctions to regulate muscle contractions, but its detection in the hemolymph of some insects suggests that it may also function as a neurohormone (37).

Interestingly, experiments in the cockroach *Blaberus craniifer* suggested a role for Proctolin in regulating yolk uptake by the oocyte (i.e., vitellogenesis) (38). The biogenic monoamine octopamine acts as a neurotransmitter, neuromodulator, and neurohormone and plays important roles in invertebrate behavior and physiology, including GSC regulation, ovulation, contraction of muscles in the reproductive tract, and movement of male and female gametes to ensure fertilization (39,40). *GABA-B-R3* encodes one of the receptors for the neurotransmitter γ- aminobutyric acid (GABA), which has important roles in female reproduction in insects and mammals (41,42). Although no role for *GABA-B-R3* has been described in *D. melanogaster* oogenesis, mutation of a different GABA receptor (encoded by *Resistant to dieldrin*, or *Rdl*) causes female infertility (43). It will be interesting to explore the potential roles of these neural signaling pathways in mediating the effects of temperature on *D. melanogaster* germline growth and survival, and oocyte quality.

Conversely, *sandman* (*sand*), *G protein* β *subunit 5* (*G*β*5*), and *CG42450*—which are implicated in dopamine signaling (36)—were among genes upregulated at 18°C and downregulated at 29°C (Fig 4B and S1 and S4 Tables). Dopamine receptors are GPCRs that transduce their signal through heterotrimeric G proteins, and Gβ5 represents one of only three Gβ proteins encoded in the *D. melanogaster* genome (44). *sand* encodes a potassium channel shown to translocate to the membrane in response to dopamine during control of *D. melanogaster* sleep homeostasis (45). One of the mammalian homologs of the protein encoded by *CG42450*, Regulators of G-protein signaling (RGS) 9, has important roles in modulating dopamine signaling (46). Additionally, *Dopamine transporter* (*DAT*) was also significantly upregulated at 18°C (but not modulated at 29°C) (S1 Table). These results suggest that dopamine signaling might be modulated by temperature in *D. melanogaster* ovaries—an intriguing possibility, especially considering our recent results showing that dopamine production in the central nervous system is required for optimal survival of vitellogenic follicles (47).

### Genes involved in synaptic transmission and ion transport or in nucleosome organization and genomic integrity are uniquely upregulated at 29°C

We next performed a similar GSEA of genes that were uniquely upregulated by at least two-fold at 29°C (S3 Table). The top 10 significantly enriched "GO Biological Processes" fell into two broad categories: I) synaptic transmission and ion transport; and II) nucleosome structure and genomic integrity (Fig 5A and S3 and S4 tables).

**Fig 5.**
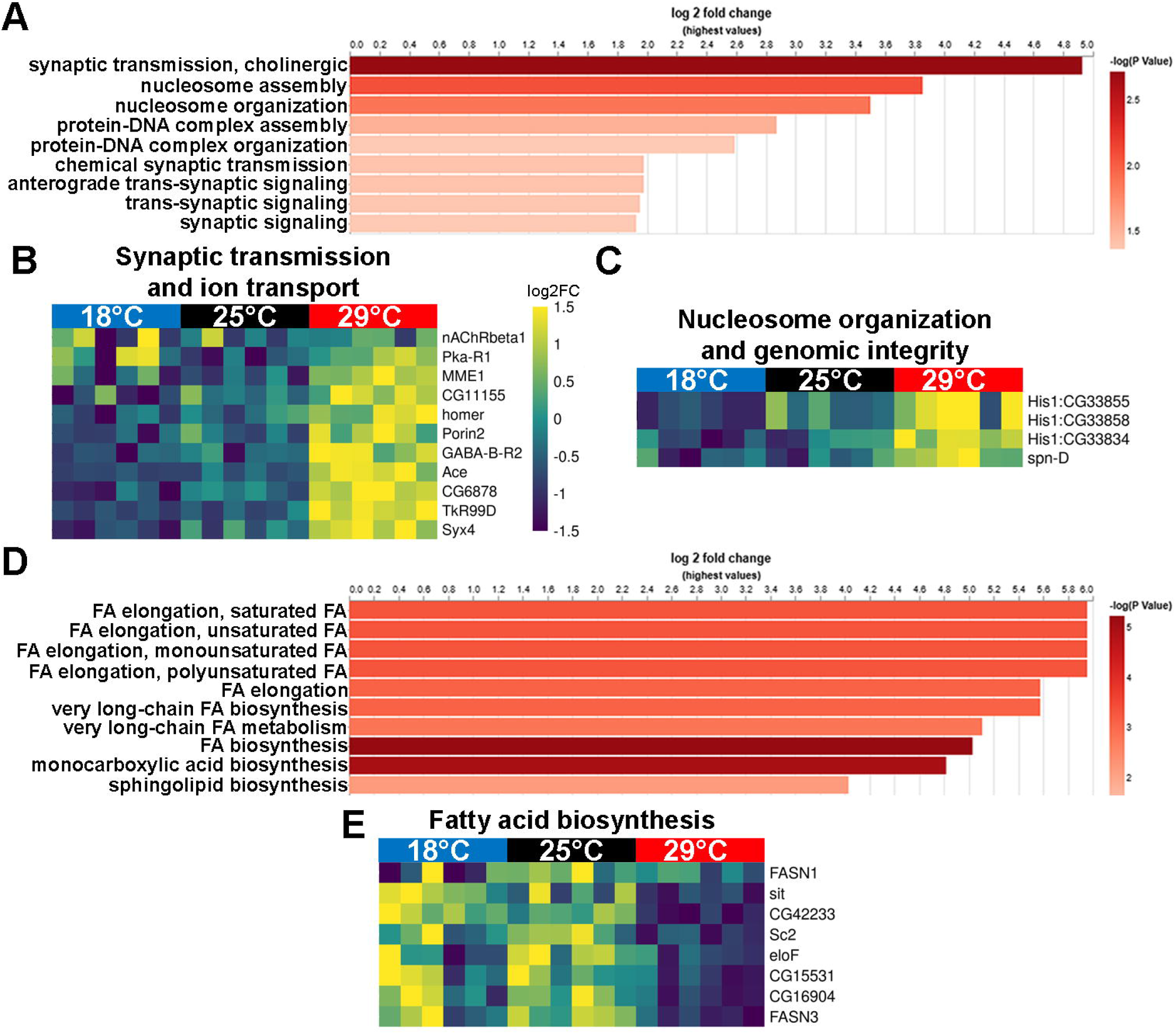
"Synaptic transmission and ion transport" and "nucleosome organization and genomic integrity" genes are uniquely upregulated, and "fatty acid biosynthesis" genes are uniquely downregulated at 29°C. (A) Bar graph showing all 9 significantly (*p*≤0.05) enriched "GO Biological Processes" among genes showing at least two-fold upregulation at 29°C and no regulation at 18°C. These enriched "GO Biological Processes" fall into "synaptic transmission and ion transport" and "nucleosome organization and genomic integrity" broad categories (S4 Table). (B and C) Heatmaps showing all 29°C upregulated genes (*q*≤0.05) involved in synaptic transmission and ion transport (B) or nucleosome organization and genomic integrity (C). (D) Bar graph showing the 10 most significantly (*p*≤0.05) enriched "GO Biological Processes" among genes showing at least two-fold downregulation at 29°C and no regulation at 18°C, which fall into the broad category of "fatty acid biosynthesis" (S4 Table). FA, fatty acid. (E) Heatmap showing all 29°C downregulated genes (*q*≤0.05) involved in fatty acid biosynthesis. Yellow and blue represent upregulated and downregulated genes, respectively.

In the first category, *GABA-B-R2* was among uniquely upregulated genes at 29°C (Fig 5B and S1 and S4 Tables), consistent with the upregulation of *GABA-B-R3* (Fig 4B). *nicotinic Acetylcholine Receptor* β*1* (*nAChR*β*1*), whose loss of function causes female sterility (48), and *Acetylcholine esterase* (*Ace*) were also upregulated at 29°C (Fig 5B and S1 and S4 Tables).

Other uniquely upregulated genes at 29°C were *homer* and *CG11155* (Fig 5B and S1 and S4 Tables). *CG11155* encodes an ionotropic glutamate receptor (49), while *homer* encodes an evolutionarily conserved adaptor for metabotropic glutamate receptors with roles in sleep regulation (50), although no roles have been described in the ovary. It is also interesting that *Tachykinin-like receptor at 99D* (*TkR99D*) and *Protein kinase, cAMP-dependent, regulatory subunit type 1* (*Pka-R1*) are upregulated at 29°C (Fig 5B and S1 and S4 Tables). Tachykinins are highly conserved neuropeptides with well-known roles in nociception and inflammation in vertebrates (51). In *D. melanogaster* larvae, *Tachykinin* and *TkR99D* are required for sensitization of sensory neurons to thermal stimulation at 45°C and 48°C following tissue damage (52). Similarly, cAMP-PKA signaling has conserved roles in mediating the effects of various extracellular stimuli during persistent pain after injury (53). Potential functions of these genes during oogenesis and/or in response to warm temperatures should be investigated.

The second category of 29°C upregulated genes included *His1:CG33855*, *His1:CG33858*, and *His1:CG33834* (Fig 5C and S1 and S4 Tables), which are members of the canonical linker histone H1 family (encoded by 23 clustered genes in *D. melanogaster*) (36). In *D. melanogaster*, canonical histone H1 has roles in heterochromatin maintenance and genome stability by promoting DNA repair and repressing TE activity (54). It is tempting to speculate that H1 genes are upregulated in response to reduced stability of heterochromatin, which has been reported at elevated temperatures (55). Another gene upregulated at 29°C, *spindle D* (*spn-D*) (Fig 5C and S1 and S4 Tables), functions in homologous recombination-mediated DNA repair during meiosis (56). Temperature is known to influence meiotic crossover frequency (57). Thus, the upregulation of *His1* and *spn-D* genes may reflect a coordinated response to temperature-induced perturbation of chromatin organization and meiotic DNA repair. Such a response could help mitigate the deleterious effects of 29°C on the germline, including increased germline death and decreased oocyte quality (13).

### 29°C uniquely downregulates fatty acid biosynthesis genes in the ovary

Strikingly, the top 10 significantly enriched "GO Biological Processes" identified by our GSEA of genes uniquely downregulated by at least two-fold at 29°C all fell into the broad category of fatty acid biosynthesis (Fig 5D and 5E and S1, S3 and S4 Tables). *elongase F* (*eloF*), *CG16904*, and *stuck in traffic* (*sit*) encode fatty acid elongases (orthologous to human ELOVL1 and ELOVL7) involved in very long-chain fatty acid biosynthesis (36). *Sc2* encodes a very long-chain enoyl-CoA reductase (orthologous to human TECR and TECRL) with roles in sphingolipid metabolism and very long-chain fatty acid biosynthesis (36). Similarly, *Fatty acid synthase 1* (*FASN1*) and *FASN3* are involved in fatty acid biosynthesis (36). Germline-specific knockdown of *FASN1* and *FASN3* in the germline reduces the number of lipid droplets in germ cells (58). Knockdown of other elongase genes or of *FASN3* in hepatocyte-like oenocytes leads to failure of egg activation and female sterility (59). Although their intrinsic roles in the germline are less clear, it is possible that downregulation of lipid biosynthesis negatively impacts the germline, contributing to the increased death of vitellogenic follicles and reduced oocyte quality at 29°C (13). *CG15531* encodes a stearoyl-CoA 9-desaturase (orthologous to human SCD and SCD5) with roles in production of saturated fatty acids (36). Human SCD5 is expressed in specific tissues (including ovaries) and functions in lipid remodeling and cell signaling (60). It has been well established (originally in bacteria) that maintenance of membrane fluidity under temperature stress is crucial and involves regulation of fatty acid saturation—e.g., higher levels of saturated fatty acids prevent excessive membrane fluidity at elevated temperatures (61); homeoviscous adaptation most likely occurs in *D. melanogaster* ovaries responding to different temperatures.

### Genes involved in muscle function and actin cytoskeleton are uniquely upregulated at 18°C

Mild cold improves GSC maintenance, slows germline development, and increases oocyte quality—i.e., the ability of oocyte to be fertilized and support successful embryonic development (13). Genes actively upregulated at 18°C against thermodynamic constraints might therefore offer interesting insight into potential mechanisms underlying the slower growth and improved germline quality in mild cold. The top 10 significantly enriched "GO Biological Processes" based on GSEA of genes that were uniquely upregulated by at least two-fold at 18°C fell into three broad categories: I) muscle function and actin cytoskeleton; II) meiotic DNA double-strand break formation; and III) reactive oxygen species metabolism (Fig 6A and S3 and S4 tables). However, the "GO Biological Processes" included in the latter two categories were not significantly enriched among all 18°C upregulated genes (S3 Table) and will not be further discussed.

**Fig 6.**
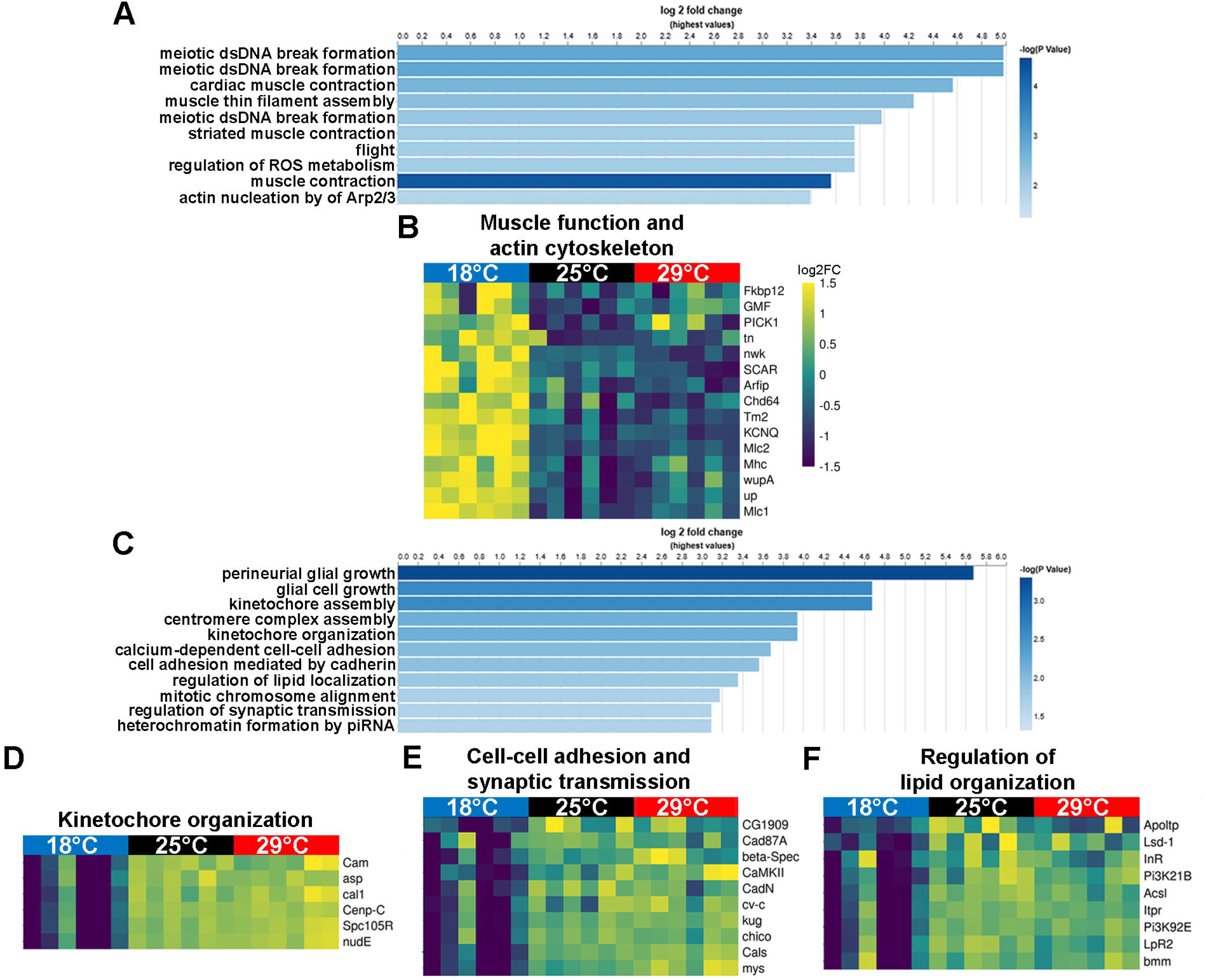
"Muscle function and actin cytoskeleton" genes are uniquely upregulated, and "kinetochore organization", "cell-cell adhesion and synaptic transmission", and "regulation of lipid organization" genes are uniquely downregulated at 18°C. (A) Bar graph showing the 10 most significantly (*p*≤0.05) enriched "GO Biological Processes" among genes showing at least two-fold upregulation at 18°C and no regulation at 29°C, which fall in the broad category of "muscle function and actin cytoskeleton" (S4 Table). (B) Heatmap showing all 18°C upregulated genes (*q*≤0.05) involved in muscle function and actin cytoskeleton. (C) Bar graph showing the 10 most significantly (*p*≤0.05) enriched "GO Biological Processes" among genes showing at least two-fold downregulation at 18°C and no regulation at 29°C, which fall into "kinetochore organization", "cell-cell adhesion and synaptic transmission", and "regulation of lipid organization" broad categories (S4 Table). (D-F) Heatmaps showing all 18°C downregulated genes (*q*≤0.05) involved in kinetochore organization (D), cell-cell adhesion and synaptic transmission (E), or regulation of lipid organization (F). Yellow and blue represent upregulated or downregulated genes, respectively.

Many genes in the "muscle function and actin organization" category encode proteins that are important for muscle contraction, which requires complex interactions between myosin and actin filaments (62). These include *Myosin heavy chain* (*Mhc*), *Myosin light chain 2* (*Mlc2*), *Myosin alkali light chain 1* (*Mlc1*), *upheld* (*up*; which encodes Troponin T), *wings up A* (*wupA*; which encodes Troponin I), *Tropomyosin 2* (*Tm2*), *thin* (*tn*; which encodes a ubiquitin ligase that regulates myofibrils), *Chd64* (which encodes a calponin), *FK506-binding protein 12kD* (*Fkbp12*), and *KCNQ potassium channel* (*KCNQ*) (Fig 6B and S3 and S4 Tables) (36). However, actomyosin complexes also have roles in non-muscle cells to promote force generation and enable cell adhesion, migration, division, and tissue biomechanics (63). During *D. melanogaster* oogenesis, the actomyosin cytoskeleton in follicle cells helps restrict the forces of the growing germline (64). Notably, contractility of actomyosin in the follicular epithelium also controls the opening of tricellular junctions (i.e., patency) to allow yolk proteins—which support later embryo development—to be taken up by the oocyte (65). *Tm2*, *wupA*, and *KCNQ* are all maternally required for early embryonic development—mutant females are sterile because the embryos they give rise to have defects during early syncytial mitotic divisions and fail to develop into larvae (66,67). Other genes upregulated at 18°C encode regulators of the actin cytoskeleton, such as *nervous wreck* (*nwk*), *Glia maturation factor* (*GMF*), and *SCAR*. *GMF* is required for robust dynamics of actin-rich protrusions in migrating border cells (68)—somatic cells required during oogenesis for morphogenesis of the eggshell structure known as the micropyle that allows sperm entry for fertilization (11). Similarly, loss of *SCAR* cell-autonomously delays the migration of border cells (69).

It is possible that at least some of these 18°C upregulated genes might contribute to high oocyte quality by acting during oogenesis to increase the success rate of key morphogenetic events (e.g., border cell migration) and yolk accumulation (for future embryo consumption) and/or by virtue of being maternally required mRNAs. It is also conceivable that these mild cold-induced genes have more indirect roles during oogenesis by acting in the mesoderm-derived muscle layers of the ovary—the epithelial sheath surrounding individual ovarioles or the peritoneal sheath, which surrounds each entire ovary—or in the oviduct muscle (70) through yet unidentified crosstalk. These possibilities should be experimentally tested in future studies.

### Genes involved in kinetochore organization, cell-cell adhesion and synaptic transmission, and regulation of lipid localization were uniquely downregulated at 18°C

The top 10 significantly enriched “GO Biological Processes” among genes uniquely downregulated by at least two-fold at 18°C fell into four broad categories: I) kinetochore organization; II) cell-cell adhesion and synaptic transmission; III) regulation of lipid localization; and IV) glial growth (Fig 6C and S3 and S4 Tables). We will not discuss the "glial growth" category because the associated "GO Biological Processes” were not significantly enriched among all 18°C downregulated genes (S4 Table).

Genes in the first category include *Centromeric protein-C* (*Cenp-C*), *Spc105-related* (*Spc105-R*), *nudE*, *chromosome alignment defect 1* (*cal1*), *Calmodulin* (*Cam*), and *abnormal spindle* (*asp*) (Fig 6D and S3 and S4 Tables). During cell division, chromosome segregation is mediated by a spindle comprising many microtubules, a subset of which attaches to the kinetochore protein complex that assembles on centromeres (71). *cal1* encodes a chaperone that incorporates the centromeric chromatin protein CENP-A into centromeric nucleosomes, *Cenp-C* encodes an inner kinetochore protein that links CENP-A to the outer kinetochore, and *Spc105-R* encodes an outer kinetochore protein that helps mediate kinetochore-microtubule association (71). *nudE* encodes a protein that associates with the kinetochore and the spindle (72), while *asp* encodes a microtubule-associated protein with a calmodulin-binding domain (73). Although *Cenp-C* and *cal1* are essential genes, their partial loss of function causes female sterility and chromosome segregation defects owing to their requirement for centromere clustering and pairing during early meiosis (74). Consistent with their critical roles in kinetochore assembly, *Cenp-C* or *cal1* knockdown leads to defects in mitotic female GSCs (75,76). In *D. melanogaster* oocytes, SPC105-R is required for stable metaphase I arrest by recruiting proteins involved in outer kinetochore assembly, cohesion, and stable end-on microtubule attachments (77). *asp* is necessary for normal germline proliferation and oocyte differentiation during oogenesis, and it is maternally provided for mitotic divisions in the early embryo (78,79). Calmodulin is a major Ca^2+^ sensor and has conserved roles in cell proliferation across eukaryotes (80). The downregulation of "kinetochore organization" genes might simply reflect the slower oogenesis at 18°C (13), although we cannot exclude other possibilities.

In the "cell adhesion and synaptic transmission" category (Fig 6E and S3 and S4 Tables), *kugelei* (*kug*, also known as *Fat2*), *Calsyntenin* (*Cals*), *Cadherin 87A* (*Cad87A*), and *Cadherin-N* (*CadN*) encode cadherin superfamily members, *myospheroid* (*mys*) encodes the β-subunit of integrin, β*-Spectrin (*β*-Spec*) encodes a component of the actin cytoskeleton, and *crossveinless c* (*cv-c*) encodes a RhoGTPase activating protein (36). Cadherins mediate homotypic cell-cell adhesions, while integrins mediate adhesion between cells and the extracellular matrix, and both stimulate the Rho family of GTPases to regulate the actin cytoskeleton (81). Mutations in *kug* or *mys* disrupt the orientation of follicle cell basal actin filaments leading to the production of abnormal, round eggs (82). Similarly, homozygous β*-Spec* mutant follicle cells have disrupted basal actin filament orientation (83). Other genes in this category include *CG1909*, which encodes the homolog of human receptor associated protein of the synapse (RAPSN); *chico*, which encodes a key substrate of the insulin receptor; and *Calcium/calmodulin-dependent protein kinase II* (*CaMKII*), which encodes a downstream effector of calmodulin with a potential role in ovulation (36). *chico* has germline-autonomous roles in stimulating GSC proliferation, follicle growth and vitellogenesis (84), suggesting that reduced insulin signaling might contribute to the lower rates of oogenesis observed at 18°C (13).

In agreement with a potential reduction in insulin signaling at 18°C, downregulated genes in the "regulation of lipid localization" category include *Insulin-like receptor* (*Inr*), *Phosphatidylinositol 3-kinase 92D* (*Pi3K92E*), and *Pi3K21B*, which encodes an adaptor protein that binds to Pi3K92E (36) (Fig 6F and S3 and S4 Tables). Indeed, activation of the insulin receptor promotes GSC division and follicle growth through Pi3K signaling (84). Insulin signaling is also known to regulate lipid metabolism (85). Accordingly, genes encoding lipid-binding proteins or enzymes involved in lipid metabolism were also downregulated, including *Apolipoprotein lipid transfer particle* (*Apoltp*), *Lipophorin receptor 2* (*LpR2*), *brummer* (*bmm*),

*Acyl-CoA synthetase long-chain* (*Acsl*), and *Lipid storage droplet 1* (*Lsd-1*). *LpR2* null females produce oocytes with severely reduced levels of neutral lipids (86). The triglyceride lipase encoded by *bmm* is required for actin remodeling in nurse cells during late oogenesis, likely through a mechanism involving arachidonic acid release from lipid droplets for prostaglandin production (87). The final gene in this category, *Inositol 1,4,5,-trisphosphate receptor* (*Itpr*), encodes an intracellular ligand gated calcium channel. Hypomorphic *Itpr* global mutants have poor fertility (88) and germline clones lacking *Itpr* function fail to yield viable eggs or embryos (89). Reduction in expression of these genes may be caused by lower insulin signaling levels and reflect the slowdown of oogenesis in mild cold.

### Chronic mild cold and warm temperature modulate different sets of transposable element transcripts

Consistent with numerous studies demonstrating that temperature influences TE expression across diverse organisms (90–92), we observed temperature-dependent modulation of TE transcripts in our ovarian transcriptomic data (Fig 1I-1L and S1 Table). A unique strength of our study, however, is the side-by-side comparison of TE expression under both mild cold and warm temperatures relative to the standard 25°C. Of the total 127 types of TEs identified in our ovarian transcriptomes (S1 Table), 102 were modulated according to temperature (S5 Table). A clustered heat map showed specific TE regulation signatures at 29°C and 18°C (Fig 7A). At 29°C, 23 TE types were upregulated and 19 downregulated, while at 18°C, 32 TE types were upregulated and 32 downregulated (Fig 7B-7E and S5 Table). The relatively large number of TEs upregulated at 18°C was unexpected, given that TE activation is commonly associated with heat stress in *Drosophila* species (93). Interestingly, genes encoding multiple proteins involved in processing of piRNAs (GO:0034587)—which have conserved roles in silencing TE expression in the germline (94)—were downregulated at 18°C but were unchanged or upregulated at 29°C (S2 Fig and S1 Table), suggesting that different mechanisms are involved in TE modulation at 29°C versus 18°C.

**Fig 7.**
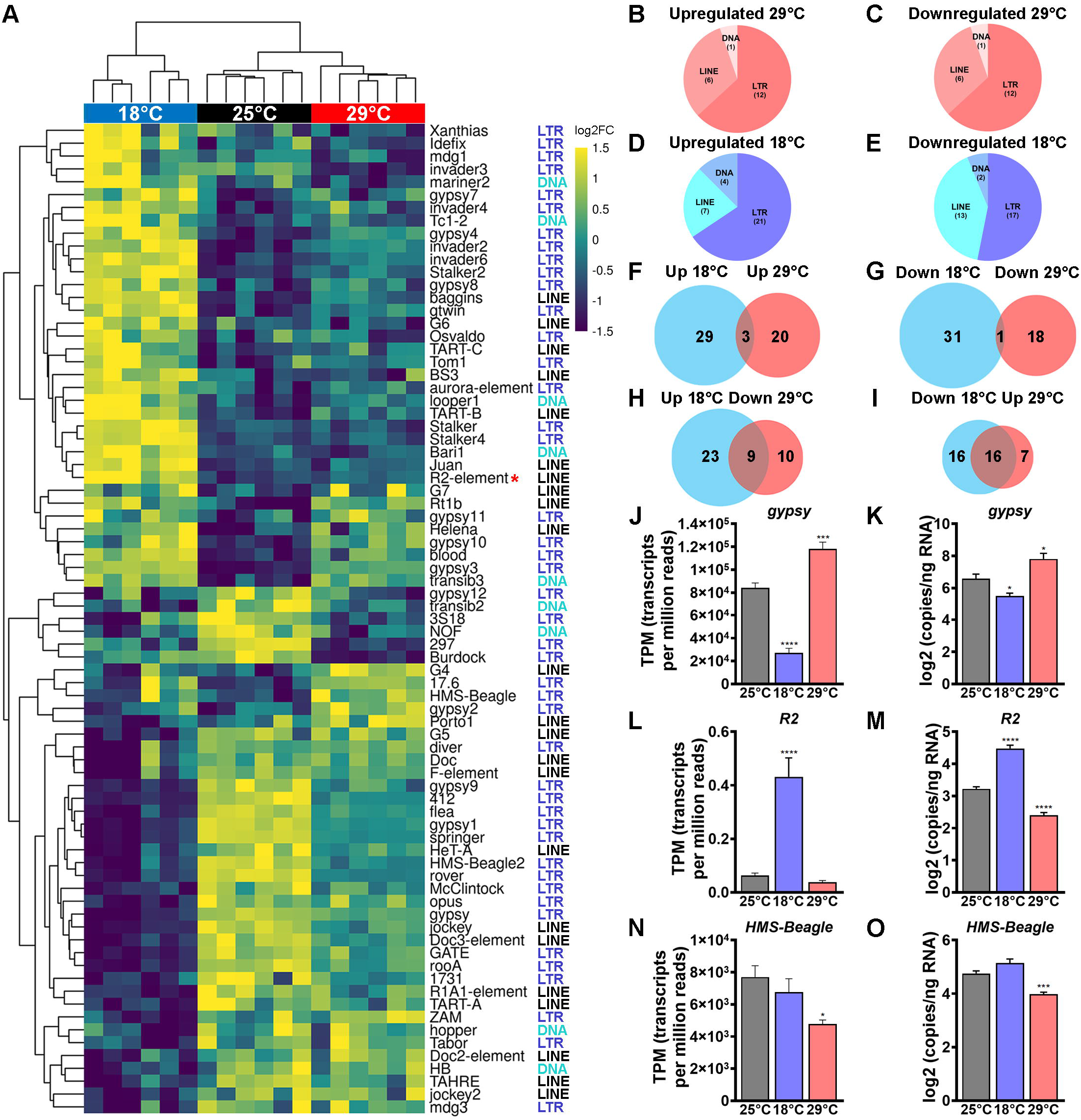
Mild cold and warm temperatures modulated specific TE transcripts. (A) Heatmap showing temperature-regulated (*q*≤0.05) TE types. Expression level is scaled per TE in each row, with yellow and blue representing upregulated and downregulated TEs, respectively. The red asterisk indicates *R2*. LTR, long terminal repeat element (Class I retrotransposon); LINE, long interspersed nuclear element (Class I retrotransposon); DNA, Class II TEs that mobilize via a DNA intermediate. (B-E) Pie charts showing the relative number of TE types belonging to different classes that are significantly (*q*≤0.05) upregulated (B) or downregulated (C) at 29°C, or upregulated (D) or downregulated (E) at 18°C. (F-I) Venn diagrams of temperature-modulated TEs, illustrating overlap among TEs regulated in the same (F and G) or opposite (H and I) directions by 18°C versus 29°C (relative to 25°C control). (J-O) Bar graphs showing expression levels of *gypsy* (J and K), *R2* (L and M), or *HMS-Beagle* (N and O) at different temperatures. (J, L, and N) show transcripts per million reads based on transcriptomics analysis (S1 Table), while (K, M, and O) show number of transcripts per ng of RNA in logarithmic scale based on nanoplate digital PCR (S5 Table). Data are shown as mean ± SEM from six (J, L, and N), or nine (K, M, and O) biological replicates. \**p*≤0.05; \*\*\**p*≤0.001; \*\*\*\**p*≤0.0001, One-way ANOVA.

TEs can affect their host genomes differently depending on their mode of transposition (94). Class I TEs, or retrotransposons, mobilize through reverse transcription followed by integration, leaving the original copy intact; these include long terminal repeat (LTR) elements and long interspersed nuclear elements (LINEs). By contract, Class II TEs mobilize via a DNA intermediate, for example, via a cut-and-paste mechanism mediated by transposases that generate double-stranded DNA breaks (94). We therefore examined whether there were differences in the classes of TEs modulated at 29°C versus 18°C. Retrotransposons are the most abundant TE class in *D. melanogaster* (93); accordingly, over 50% of TEs modulated by both temperatures in the ovary were LTRs, followed by LINEs, and, lastly, class II DNA TEs (Fig 7A-7E). However, the relative proportions of modulated TE families differed at 29°C versus 18°C. At 29°C, upregulated TE types were 74% LTRs, 17% LINEs, and 1% DNA TEs, and downregulated TE types were 63% LTRs, 32% LINEs, and 1% DNA TEs. At 18°C, 66% LTRs, 22% LINEs, and 13% DNA TEs were upregulated, while 53% LTRs, 41% LINEs, and 6% DNA TEs were downregulated (Fig 7B-7E). These results suggest that mild cold disproportionally modulates class II DNA TEs and differentially regulates class I TEs, with relatively more LINEs and fewer LTRs being modulated at 18°C compared to 29°C. The bias toward DNA TE upregulation at 18°C in the ovaries appears counterintuitive—DNA TE transposases are known to actively generate double-stranded DNA breaks (94) and yet the quality of the female germline is better maintained at 18°C compared to 25°C or 29°C (13).

To further compare the TE temperature response patterns, we generated Venn diagrams to visualize TE types regulated in opposite directions, in the same direction, or uniquely modulated at 18°C or 29°C (Fig 7F-7I and S5 Table). Of all 102 temperature-modulated TE types, only three (3%)—*Burdock*, *297*, and *invader3*—were upregulated at both 18°C and 29°C (Fig 7F and S5 Table), while only *G4* (1%) was downregulated at both temperatures (Fig 7G and S5 Table), indicating that suboptimal temperatures have minimal nonspecific effects on TE expression. By contrast, 25 TE types (25%) were modulated in opposite directions at 18°C versus 29°C, while the remaining 52 (52%) were uniquely modulated by either mild cold or warm temperature (Fig 7H and 7I). Among TE types with opposite expression patterns at 18°C and 29°C, nine were downregulated at 29°C and upregulated at 18°C—*TART-B*, *Stalker2*, *gypsy4*, *blood*, *invader2*, *baggins*, *gtwin*, *invader6*, and *gypsy3*—while 16 were upregulated at 29°C and downregulated at 18°C—*rover*, *gypsy9*, *HMS-Beagle2*, *flea*, *412*, *springer*, *gypsy1*, *jockey*, *gypsy*, *rooA*, *McClintock*, *Doc3-element*, *HeT-A*, *R1A1*, *1731*, *GATE*) (Fig 7H-7K and S5 Table). Among uniquely modulated TEs, 20 were upregulated and 15 were downregulated at 18°C, while 4 were upregulated and 6 downregulated at 29°C (Fig 7F-7I). For example, *R2* was upregulated at 18°C and unchanged (based on transcriptomics; Fig 7L) or downregulated (based on nanoplate digital PCR; Fig 7M) at 29°C and *HMS-Beagle* was downregulated at 29°C and unchanged at 18°C (Fig 7N and 7O). Our results agree with two previous studies showing that TE expression can be modulated by suboptimal temperatures in *D. melanogaster* (24,95).

Importantly, our analysis reveals a high degree of specificity in how TE expression is modulated by warm temperature versus mild cold (relative to the control 25°C temperature) in *D. melanogaster* ovaries. Although TE activation is often associated with DNA damage, TEs can have beneficial roles (96,97). Additional studies are needed to determine whether individual temperature-modulated TEs play positive, negative, or neutral roles in the ovarian response to mild cold or warm temperature. Future research should also examine a potential connection between the differential regulation of TE types at different temperatures and variations in the abundance of certain TE types in *Drosophila* populations found in different geographical and environmental contexts (98,99).

### Mild cold specifically downregulates genes involved in chromatin regulation

Precise regulation of TE transcription involves the dynamic regulation of chromatin structure and, conversely, TEs can also control the spreading of repressive chromatin marks to neighboring genomic regions (94). Given that 29°C versus 18°C differentially modulates TE expression in the ovary (Fig 7), we wondered if there might also be temperature-specific changes in genes involved in chromatin regulation. Our GSEA results revealed significantly enriched “GO Biological Processes” related to chromatin regulation among genes uniquely downregulated at 18°C—although not among 18°C upregulated or 29°C modulated genes (S3 Table). These significantly enriched GO terms—GO:0140966 (piRNA-mediated heterochromatin formation), GO:0141006 (Transposable element silencing by piRNA-mediated heterochromatin formation), GO:0031048 (Regulatory ncRNA-mediated heterochromatin formation), and GO:0141005 (Transposable element silencing by heterochromatin formation) (S3 Table)— encompassed *small ovary* (*sov*) and *windei* (*wde*), which were downregulated by at least two-fold, and *Suppressor of variegation 3-3* [*Su(var)3-3*], *Su(var)2-10*, *MEP-1*, *asterix* (*arx*), *eggless* (*egg*), *ADD domain-containing protein 1* (*ADD1*), *Argonaute 2* (*AGO2*), *CG5877*, *SET domain containing 2* (*Set2*), *winged eye* (*wge*), *Nuclear receptor binding SET domain protein* (*NSD*), and *absent, small, or homeotic discs 1* (*ash1*), downregulated to a lesser extent at 18°C (Fig 8A and S1 Table).

**Fig 8.**
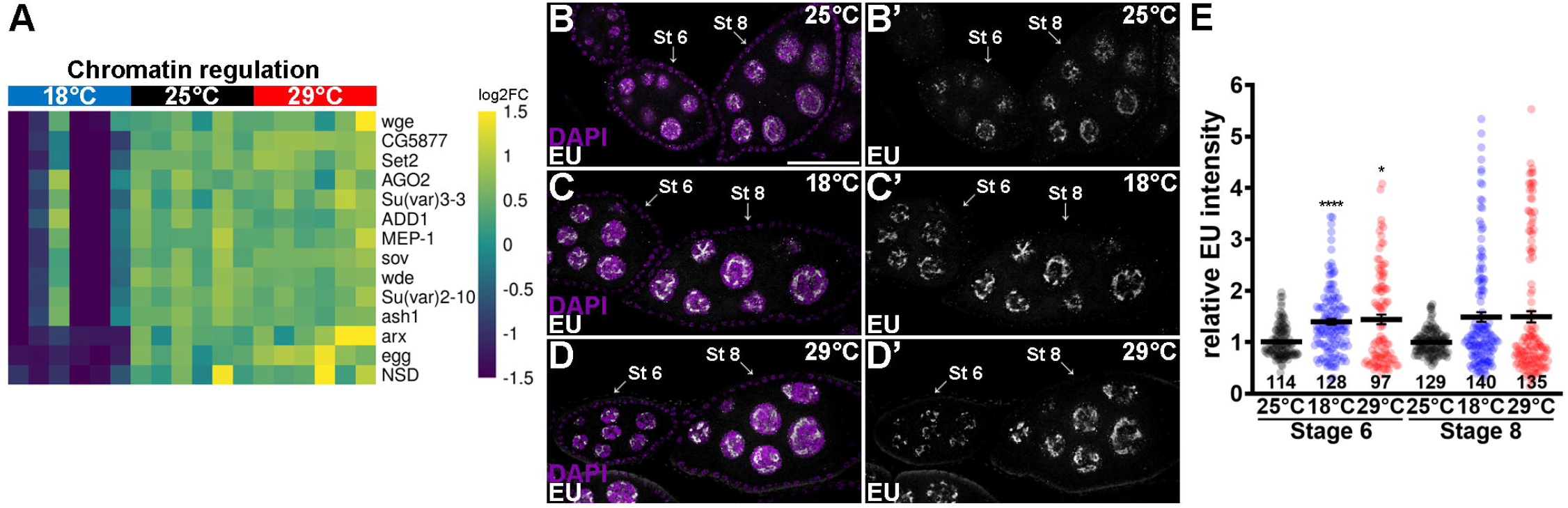
Mild cold downregulates "chromatin regulation" genes. (A) Heatmap showing genes involved in chromatin regulation that are significantly (*q*≤0.05) downregulated by mild cold. Yellow and blue represent upregulated and downregulated genes, respectively. (B-D’) Single optical slices of part of ovarioles from females maintained at 25°C control (B), 18°C (C), or 29°C (D) for five days. Stage 6 (St 6) and stage 8 (St 8) follicles are indicated by arrows. EU (white) labels newly transcribed RNA; DAPI (magenta) labels nuclei. Single channel for EU is shown in (B’, C’, and D’). Scale bar: 50 µm. (E) Dot plot showing quantification of EU intensity normalized by DAPI intensity in stage 6 and 8 nurse cell nuclei with 25°C control set as 1. Lines indicate mean ± SEM from two biological replicates. The numbers of nurse cells analyzed are shown at the bottom of graph. Numbers of follicles analyzed: 36 stage 6 and 40 stage 8 at 25°C; 37 stage 6 and 41 stage 8 at 18°C; 32 stage 6 and 41 stage 8 at 29°C. \**p*≤0.05; \*\*\*\**p*≤0.0001, Kruskal- Wallis test.

Many of the proteins encoded by these downregulated genes are involved in the process through which piRNA-loaded Piwi protein targets nascent TE transcripts for deposition of the heterochromatic repressive mark H3K9me3 in the germline (94). The zinc finger protein Arx (also known as Gstf1), the SUMO E3-ligase encoded by *Su(var)2-10*, and the MEP-1 Kru ppel- type zinc-finger protein interact with Piwi, with Su(var)2-10 linking the Piwi complex to the histone H3K9 methyltransferase Egg (also known as SetDB1) and its cofactor Wde—all of which are required for H3K9 methylation and TE silencing in the *D. melanogaster* germline (94,100–103). Follicles containing *egg* or *wde* mutant germline cysts do not progress past early pre-vitellogenic stages (103,104). ADD1 has complex roles in chromatin regulation and interacts with Egg and the heterochromatin protein HP1a (105). Su(var)3-3 (also known as Lsd1) is a lysine-specific demethylase for H3K4 and H3K9 that regulates developmental genes and is required for Piwi-dependent TE silencing during oogenesis (106). Sov is a multi-zinc finger protein important for general heterochromatin formation that is recruited to TE sites in a Piwi- dependent manner and is required in the female germline for TE silencing, genome integrity, maintenance of GSCs, and production of high-quality oocytes (107,108). Overall, the downregulation of genes encoding these proteins agrees with the downregulation of piRNA processing genes and upregulation of a subset of transposons at 18°C (Fig 7A and S2 Fig and S1 Table).

Other 18°C downregulated genes are known to function in various ways to modulate chromatin and silence TEs. AGO2 is an Argonaute/Piwi family protein involved in processing small interfering RNAs, microRNAs, and piRNAs to regulate chromatin structure and repress TE expression (109,110). *CG5877* is the ortholog of *C. elegans nrde-2* and human *NRDE2*, which encode an RNA splicing factor required for nuclear RNA interference downstream of AGO (111,112). *wge* controls histone transcription and is required for TE repression and genome integrity (113). The *Ash1*, *NSD*, and *Set2* encode H3K36-specific methyl transferases required for TE repression in the brain (114). *Set2* is also required in the female germline for differentiation (115), while global *ash1* hypomorphic mutant females are fertile at 20°C but stop producing eggs within 2 days at 27°C (116), suggesting a stronger requirement at higher temperatures.

The downregulation of multiple genes involved in heterochromatin formation and TE silencing suggests that ovaries at 18°C might exhibit a more transcriptionally active state overall. We therefore incubated freshly dissected ovaries from females maintained at 18°C, 29°C, or control 25°C temperatures with 5-ethynyl uridine (EU) to label newly synthesized RNA and assess global transcription in developing follicles (Fig 8B-8D’). Interestingly, the intensity of EU labeling was higher not only at 18°C but also at 29°C compared to 25°C (Fig 8E).

Acetylation of H3K9—which is associated with transcriptionally active chromatin (117)—was consistently increased at 18°C but not at 29°C (S3A-S3C’ Fig). We also observed a slight increase in repressive H3K9me3 at 18°C (S3D-S3F’ Fig), while HP1a was more robustly increased at 18°C (S3G-S3I’ Fig). Although HP1a is a well-known heterochromatin protein involved in TE repression and binding to methylated H3K9 (118), it is also required for positive regulation of a subset of euchromatic genes (119,120). Understanding how chromatin regulation impacts differentially regulated TEs and protein-coding genes at 18°C and 29°C (compared to 25°C) will require genome-wide profiling using chromatin immunoprecipitation sequencing (121) to define the association of various chromatin marks with specific modulated gene loci at different temperatures.

### Larger nucleoli and elevated phosphorylated Mad levels underlie improved germline stem cell maintenance at 18°C

We were particularly intrigued by the observation that *R2* was among the most upregulated TE types in whole ovaries at 18°C (Fig 7A, 7L, and 7M and S5 Table). In many animal taxa, *R2* exclusively inserts in the 28S rRNA gene within rDNA repeat units, with *D. melanogaster* having *R2* in 10-20% of their rDNA repeats (122). In *D. melanogaster*, *R2* is required in the male germline for GSC maintenance and for maintenance of rDNA copy number through a mechanism involving *R2*-induced double-stranded breaks (123). We previously showed that *D. melanogaster* females at 18°C maintain higher numbers of GSCs that produce better quality oocytes over time compared to GSCs at 29°C or 25°C (13)—similar to findings in *C. elegans*, where cold temperature delays the exhaustion of GSCs during adulthood (124). We therefore wondered if *R2* upregulation might be occurring in GSCs of *D. melanogaster* females maintained at 18°C. Using RNA fluorescence *in situ* hybridization, we found that GSCs have increased *R2* expression at 18°C compared to either 25°C or 29°C (Fig 9A-9D). Accordingly, we also detected increased levels of double-stranded DNA breaks in GSCs at 18°C (Fig 9E-9H). *R2* expression was recently shown to be repressed by insulin signaling in male GSCs (125). Future experiments should address whether there is a mechanistic connection between the downregulation of insulin pathway components *InR*, *chico*, *Pi3K92E*, and *Pi3K21B* (Fig 6E and 6F and S4 Table) and the upregulation of *R2* at 18°C (Fig 7A, 7L, and 7M and S5 Table).

**Fig 9.**
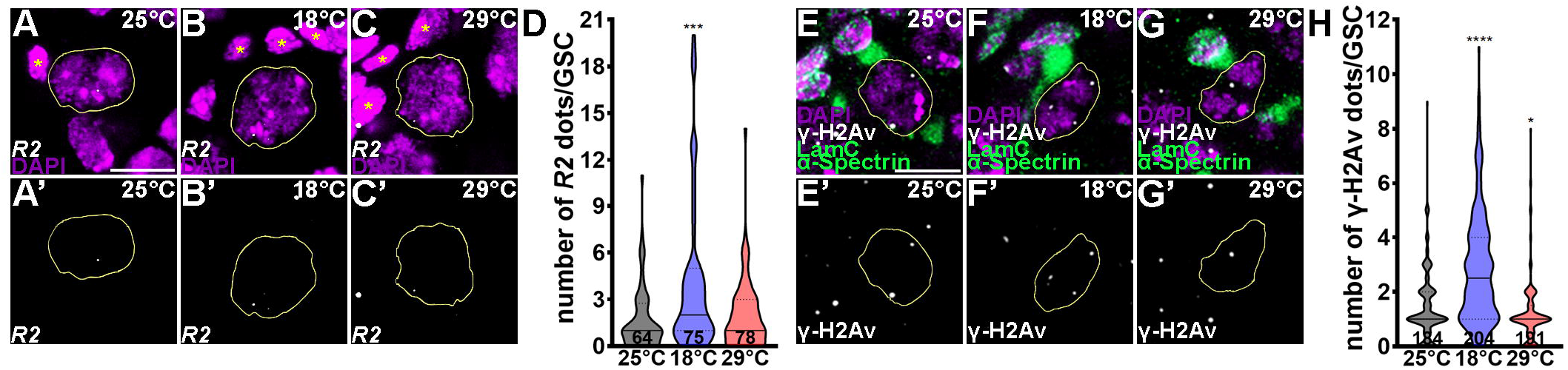
Mild cold increases *R2* transcripts and double-stranded DNA breaks in germline stem cells. (A-C’) Maximal projection images of three 1-µm optical slices of GSCs from females maintained at 25°C control (A), 18°C (B), or 29°C (C) for five days. GSC nuclei are outlined; cap cells are indicated by asterisks. *R2* Stellaris probe set (white) labels *R2* transcripts; DAPI (magenta) labels nuclei. Single channel for *R2* is shown in (A’, B’, and C’). (D) Violin plot showing quantification of the number of *R2* foci per GSC nucleus from two biological replicates. (E-G’) Maximal projection images of three 1-µm optical slices of GSCs as in (A-C). γ-H2Av (white) labels double-stranded breaks; Lamin C (LamC; green) labels cap cell nuclear envelopes; α-Spectrin (green) labels fusomes. Single channel for γ-H2Av is shown in (E’, F’, and G’). Scale bars: 5 µm. (H) Violin plot showing quantification of the number of γ-H2Av foci per GSC nucleus from three biological replicates. The numbers of GSCs analyzed are shown at the bottom of the graphs. \**p*≤0.05; \*\*\**p*≤0.001; \*\*\*\**p*≤0.0001, Kruskal-Wallis test.

*R2* is co-transcribed with pre-rRNAs by RNA polymerase I (Pol I) in the nucleolus (122), raising the possibility that *R2* upregulation in female GSCs at 18°C might reflect increased rRNA production and ribosome biogenesis. Ribosome production is positively associated with nucleolus size (126), and our ovary transcriptomics showed increased levels of *fibrillarin*, which encodes a highly conserved rRNA 2’-O-methyltransferase (127), at 18°C compared to 25°C or 29°C (S1 Table). We therefore labeled the nucleoli of GSCs from females maintained at different temperatures using antibodies against Fibrillarin and Under-developed (Udd)—a Pol I regulator (128) (Fig 10A-10C"). Both the volume and total intensity of Fibrillarin and Udd staining were significantly increased in GSCs at 18°C, compared to either 25°C or 29°C (Fig 10D-10G), indicating that nucleoli are significantly larger in GSCs from females exposed to mild cold. A recent study showed that HP1a and b proteins are required for ribosomal RNA biosynthesis and nucleolar integrity in undifferentiated mouse embryonic stem cells (129), and rDNA repeats with *R2* insertions are enriched with HP1a in *D. melanogaster* ovaries (130). In accordance, HP1a levels were also elevated in GSCs 18°C (Fig 10H-10K), similarly to what we observe in the later germline (S3G-S3I’ Fig). Taken together, these results suggest that ribosome assembly is elevated in GSCs of females exposed to mild cold.

**Fig 10.**
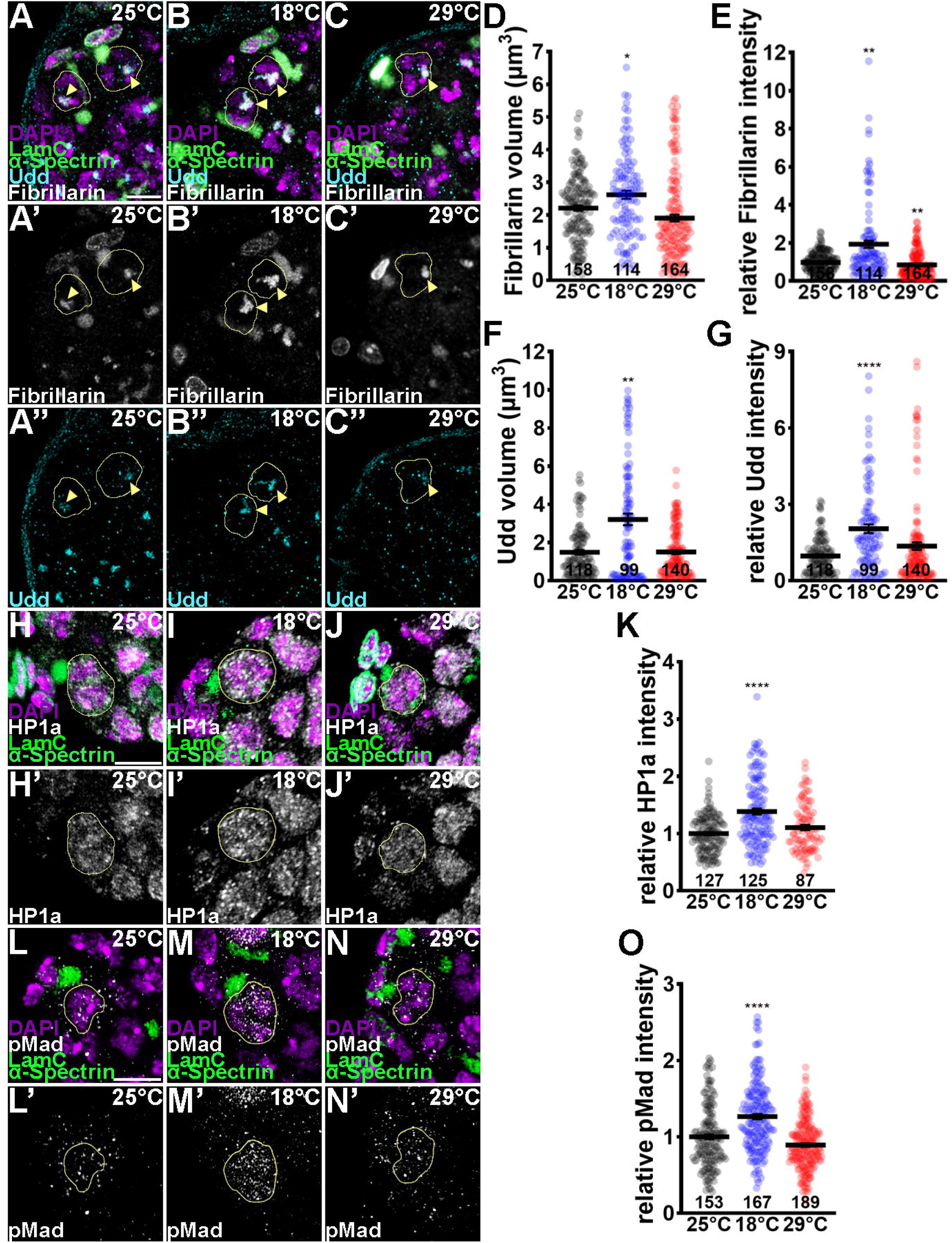
Germline stem cells have larger nucleoli and increased levels of the stemness factor pMad at 18°C. (A-C") Maximal projection images of three 1-µm optical slices of GSCs from females maintained at 25°C control (A), 18°C (B), or 29°C (C) for five days. GSC nuclei are outlined. Arrowheads indicate nucleoli. Fibrillarin (white) and Udd (cyan) label nucleoli; DAPI (magenta) labels nuclei; Lamin C (LamC) labels cap cell nuclear envelopes (green); α-Spectrin labels fusomes (green). Single channels for Fibrillarin (A’-C’) and Udd (A"-C") are shown. LamC and α-spectrin signals are visible in the fibrillarin channel because all three were detected using mouse antibodies (see Methods). (D-G) Dot plots showing quantification of fibrillarin (D and E) or Udd (F and G) volume (D and F) or relative intensity with control set as 1 (E and G). (H-J’) Maximal projection images of three 1-µm optical slices of GSCs as in (A-C). HP1a (white) single channel is shown in (H’-J’). (K) Dot plot showing relative intensity of HP1a normalized by DAPI intensity with 25°C control set as 1. (L-N’) Single optical slices of GSCs as in (A-C). pMad (white) indicates active BMP signaling, with single channel shown in (L’, M’, and N’). Scale bars: 5 µm. (O) Dot plot showing relative average intensity of pMad with 25°C control set as 1. Lines indicate mean ± SEM from two (D-G), four (K), or three (O) biological replicates. \**p*≤0.05; \*\**p*≤0.01; \*\*\*\**p*≤0.0001, One-way ANOVA (D) or Kruskal-Wallis test (E-G, K and O).

Multiple studies indicate that high levels of ribosome biogenesis are important for GSC maintenance (131), at least in part by promoting Mad expression and BMP signaling (128). Consistent with their larger nucleoli, we found that GSCs at 18°C exhibit higher pMad levels compared to those at 25°C or 29°C (Fig 10L-10O), likely contributing to the improved GSC maintenance in females maintained at 18°C (13). Altogether, our results support a working model that mild cold upregulates rRNA production and ribosome biogenesis, leading to increased levels of BMP signaling in GSCs that contribute to better GSC maintenance over time. Nucleolus assembly *in vivo* is governed by both active and thermodynamic processes (132), suggesting that the formation of larger nucleoli (and increased ribosome biogenesis) at 18°C is driven by complex mechanisms that remain to be elucidated. It is also unclear whether *R2* upregulation is merely a consequence of increased rRNA transcription or instead plays a more direct role in maintaining female GSCs at 18°C.

## Conclusion

Overall, our comprehensive transcriptomic analysis reveals complex molecular responses of *D. melanogaster* oogenesis to suboptimal temperatures and provides new insight into why germline quality improves under mild cold conditions but rapidly declines at warm temperatures. We first demonstrate that the ovary actively responds to both mild cold and warm temperatures by regulating specific sets of transcripts, indicating that these responses are not simply passive consequences of thermodynamic effects. Notably, 77% of temperature-modulated genes exhibited unique responses to individual temperatures, whereas only 21% showed reciprocal regulation between mild cold and warm conditions, further supporting the existence of distinct regulatory programs that govern the response to each temperature. For example, genes involved in neuronal signaling were enriched among reciprocally regulated genes. Intriguingly, distinct HSPs were upregulated in response to mild cold versus warm temperatures, suggesting that individual chaperones may have specialized, temperature-dependent functions. In addition, biological pathways enriched among genes selectively modulated at each temperature point to distinct mechanisms underlying adaptation to mild cold versus warm conditions (Fig 11A). For instance, upregulation of genes involved in nucleosome organization and genome integrity at 29°C may represent a protective response to chromatin instability under elevated temperatures, whereas downregulation of fatty acid biosynthesis genes at 29°C and lipid localization genes, including insulin pathway components, at 18°C may reflect temperature-specific metabolic adaptations. At 18°C, upregulation of actin cytoskeleton genes may support essential cellular processes during oogenesis and thereby contribute to improved oocyte quality. Similarly, the distinct sets of TEs upregulated or downregulated at 18°C and 29°C suggest that mild cold and warm temperatures induce different chromatin states and mechanisms of TE regulation.

**Fig 11.**
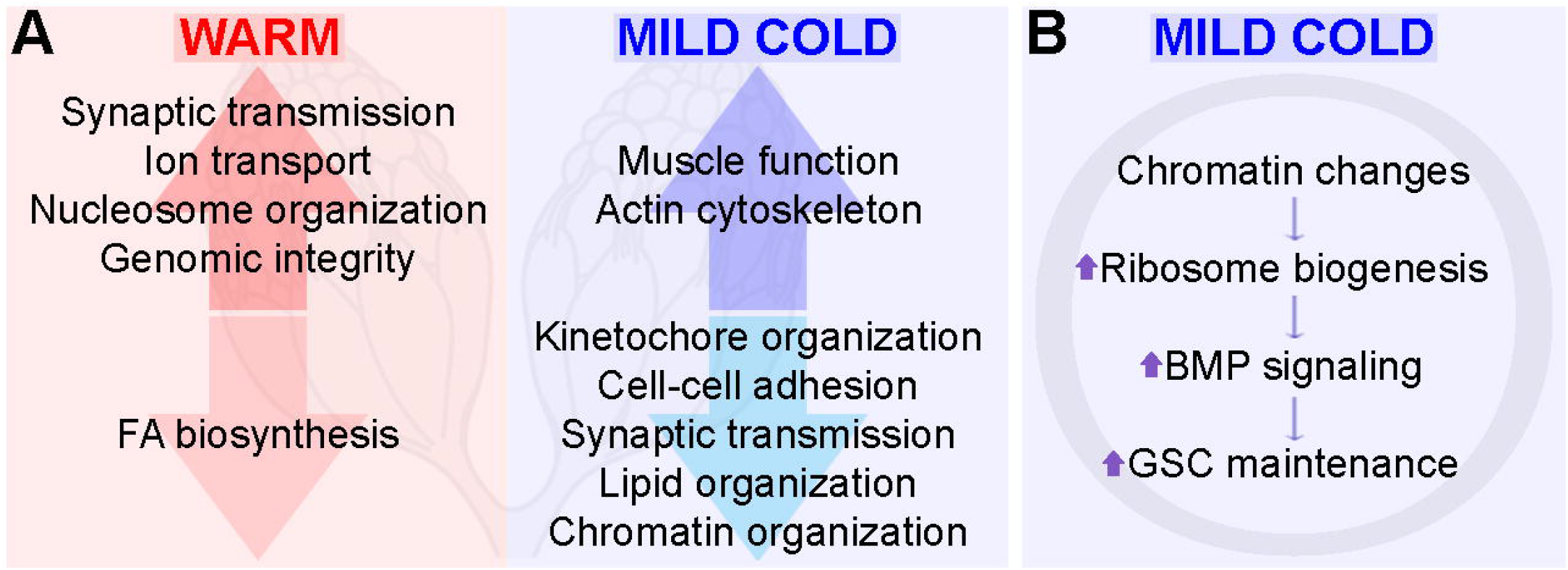
Model of molecular responses to mild cold and warm temperatures during *Drosophila* oogenesis. (A) Diagram illustrating that the ovary actively responds to mild cold and warm temperatures by selectively modulating thousands of transcripts, most of which exhibit unique temperature-dependent responses. (B) Working model proposing that mild cold induces chromatin changes and increases rRNA production thereby promoting ribosome biogenesis and elevated BMP signaling in GSCs to improve their maintenance over time.

Finally, our findings identify increased ribosome biogenesis, potentially associated with upregulation of the retrotransposon *R2*, together with enhanced BMP signaling at 18°C as potential mechanisms underlying the improved maintenance of GSCs in females exposed to mild cold (Fig. 11B). These findings raise the possibility that mild cold exposure may promote stem cell function more broadly by stimulating ribosome biogenesis. Supporting this possibility, a recent study in cultured mammalian cells demonstrated that forced activation of rRNA transcription increases nucleolar size, enhances cellular translation, and promotes neural stem cell self-renewal (133), although the study did not examine the effects of temperature on these processes. Together, our findings provide a framework for understanding how temperature-specific transcriptional and chromatin responses shape germline maintenance and quality, with broader implications for thermal biology, stem cell biology, and reproductive biology.

## Materials and Methods

### *Drosophila melanogaster* culture and experimental conditions

*y^1^ ac^1^ w^1118^* (commonly referred as *y w*) fly stocks were maintained at 21-23°C on standard medium containing 5.57% w/v cornmeal, 5.85% v/v molasses, 2.1% w/v yeast and 1.11% w/v agar. For transcriptomics sample preparation, newly eclosed flies were maintained on vials containing standard medium supplemented with dry yeast (changed daily) at 18°C (±0.5°C), 25°C (±0.5°C), or 29°C (±0.5°C) for five days at ≥ 80% humidity and 12:12 photoperiod (Darwin Chambers, IN034-LT-MP) prior to dissection. Samples were prepared in six biological replicates per temperature, with each biological replicate including 20 pairs of ovaries (Fig 1D). Each vial housed 10 couples, and a single pair of ovaries was dissected at a time and immediately snap frozen inside a 2-ml Eppendorf Safety Lock tube (containing one 5-mm bead) on dry ice. For other experiments, zero-to-one-day-old females (with males) were incubated at different temperatures as described above.

### Transcriptomics sample preparation

The University of Wisconsin–Madison Biotechnology Gene Expression Center Core (Research Resource Identifier SCR_017757) performed RNA Extraction, quality control, sequencing, and primary bioinformatic analyses of the raw data. The Stranded RNA Library was produced using the platform Illumina (NovaSeq), with a Flow Cell of 10B Full Lane and a Read Length of 2x150 (Paired-End Sequencing). In summary, 300 µl TriReagent were added to frozen ovaries in 2-ml Eppendorf Safety Lock tubes with one 5 mm bead on dry ice, homogenized for 2 minutes at 20 Hz followed by incubation on ice for 5 minutes. Then, 700 µl of TriReagent were added, followed by the addition of 200 µl of chloroform, and samples were vortexed for 15 seconds. After incubation for 2 minutes at room temperature, organic extraction followed with clean up on Qiagen RNeasy Mini Kit (lot #175010050) was performed before the on-column DNase treatment (lot #172048647). Samples were eluted with 30 µl nuclease-free water added to the column. All RNA samples were stored at -80°C and concentrations were measured on NanoDrop One Spectrophotometer and Agilent 4200 Tapestation to assess yield, purity, and integrity. The read- and mapping-level quality control metrics were of high quality and consistent with good total RNA extraction and library creation (S1 File). RNA sequencing produced an average of 38,389,394 reads across the six replicates, ranging from 35.1 to 43.3 million reads per sample, of which an average of 96% were mapped to the *D. melanogaster* genome; 60,000 reads were randomly subsampled to reduce data complexity (134). There was some slight 3’- end transcript degradation but still within acceptable levels and the pattern and duplication levels were consistent across all samples and within acceptable ranges for mRNA-Seq (S1 File). The library normalization procedure completed successfully with no outliers detected (S1 File). Part of preparing expression data for differential gene expression analysis is to determine which genes have sufficiently large counts to be retained in a statistical analysis through filtering, removing genes that have too few reads to establish any worthwhile significant expression differences (false positives) (134). Based on the analysis of those reads, 11,864 transcripts were identified (S1 Table).

### Bioinformatics analyses

Obtained sequence reads (FastQ files) were analyzed with local Galaxy installation. First, the raw reads were exposed to quality control checking, then the adaptor was trimmed by using Trimmomatic (v0.2.2b) (135). All the fragmented reads were mapped to the iso-1 reference genome (https://www.ensembl.org/genome/GCA_000001215.4) from UCSC (https://genome.ucsc.edu/index.html) by using the HISAT2 tool (136) and then we assembled the alignments into full-length transcript by using the StringTie tool.17 (137). The differential expression analysis was performed using edgeR (v.3.34.0) with a generalized linear model (138). For comparison of the experimental groups, the contrasts were 18°C versus 25°C and 29°C versus 25°C. The adjusted *p*-value (*q*-value) used to determine the cutoff for differentially expressed genes was obtained by conversion of raw *p*-values through the Benjamini-Hochberg procedure to control the overall false discovery rate (FDR) at 5% (*q*-value≤0.05) (139). For quantifying TE family expression, we obtained the *Drosophila* TE consensus library from the Bergman Lab GitHub repository (v10.2, https://github.com/bergmanlab/drosophila-transposons). We mapped the trimmed reads with bowtie2 (140) and performed the family-level transcript quantification using the TEcount.py script from TEtools (99) with default parameters. The differential TE expression analysis was performed using DESeq2 (v.1.50.2) (141). The TE family count matrices are also regularized log-transformed. For pairwise comparisons, contrasts were extracted for 18°C vs 25°C and 29°C vs 25°C. TE families with q_value ≤ 0.05 were considered statistically significant. For generation of heatmaps and PCA graphs using GraphBio (142), the expected read counts were normalized and transformed with a regularized logarithm (rlog) function that transforms the expected count data to a log2 scale (S1 Table) in a way that minimizes differences between samples, especially for genes with small counts, and that normalizes with respect to library size. Volcano plots were generated using ggplot2 (v.4.0.3) (143), and Venn diagrams were generated using DeepVenn (144). For all other graphs, we used GraphPad Prism version 10. For GSEA, genes that were significantly modulated or modulated by at least two-fold were analyzed using PANGEA with a *p*-value ≤ 0.05 (35).

### Nanoplate digital PCR (dPCR)

Six pairs of ovaries per sample were dissected and incubated in RNAlater Stabilization Solution (Thermo Fisher Scientific, AM7021) on ice. After RNAlater removal and addition of 250 μl lysis buffer from the RNAqueous-4PCR Total RNA Isolation kit (Thermo Fisher Scientific, AM1914), samples were homogenized using a motorized pestle. RNA extraction proceeded according to the manufacturer’s instructions. Complementary DNA (cDNA) was synthesized from 0.5 mg total RNA using oligo (dT) primers and SuperScript IV Reverse Transcriptase (Thermo Fisher Scientific, 18090010) according to the manufacturer’s instructions. To verify cDNA quality prior to proceeding to nanoplate digital PCR, cDNA was amplified through a 35-cycle reaction (94°C for 30 seconds, 56°C for 30 seconds and 72°C for 30 seconds) using primers for *RpL32* (5’- CAGTCGGATCGATATGCTAAGC-3’ and 5’-AATCTCCTTGCGCTTCTTGG-3’). Transcripts copy numbers for *R2*, *gypsy*, and *HMS-Beagle* were determined using nanoplate digital PCR with QIAcuity EvaGreen PCR kit (Qiagen, 250111) for all dPCR reactions (145) using the following primers: *5’-AACAGGAGAGAAAGCAGAAG-3’* (forward) and *5’-TATGCTTATCCCAGTTACGC-3’* (reverse) for *R2*; *5’-CCATACCATTTAGCCGATCA-3’* (forward) and *5’- TCTGTAGTTATGTGCCCAAC-3’* (reverse) for *gypsy*; *5’-AAATCCGTCTCCCTTAATCG-3’* (forward) and *5’-AGTGGAGTATTACGTCTGGA-3’* (reverse) for *HMS-Beagle*. cDNA samples were transferred to an 8,500-partition 96-well QIAcuity nanoplate and loaded to a QIAcuity One system (2 minutes at 95°C followed by 40 cycles of 95°C for 15 seconds, 55°C for 15 seconds and 72°C for 15 seconds). The nanoplate was imaged with an exposure duration of 100 milliseconds and gain of 2. Images were analyzed with the QIAcuity Software Suite, and the concentration of transcripts were calculated using Poisson statistical methods by the QIAcuity Software Suite. See raw data in S5 Table.

### Tissue immunostaining and fluorescence microscopy

Ovaries were dissected in Grace’s Insect Medium (Gibco, 11595-030), fixed for 15 minutes at room temperature in fixing solution [5.3% formaldehyde (Thermo Scientific, 28908) in Grace’s medium], rinsed once and washed three times for 15 minutes each in PBST [PBS plus 0.1% Triton X-100 (Sigma, T8787)]. After incubation in blocking solution [5% normal goat serum (MP Biochemicals, 642921) plus 5% bovine serum albumin (Sigma, A6003) in PBST] for at least 3 hours at room temperature or overnight at 4°C, ovaries were incubated overnight at 4°C in the following primary antibodies diluted in blocking solution: 1:400 rabbit anti-γ-H2Av phosphoS137 (Rockland #600-401-914); 1:20 mouse anti-α-Spectrin (Developmental Studies Hybridoma Bank (DSHB) 3A9; deposited by D. Branton and R. Dubreuil); 1:100 mouse anti-Lamin C (DSHB LC28.26; deposited by P. A. Fisher); 1:500 rabbit H3K9ac (Abcam #ab4441); 1:1000 rabbit H3K9me3 (Active Motif #39062); 1:800 guinea pig anti-Udd (a gift from M. Buszczak) (128).

Ovaries were rinsed and washed as above and incubated for two hours at room temperature in blocking solution containing 1:400 Alexa Fluor 488-, 568- or 633-conjugated secondary antibodies (Thermo Fisher Scientific, A10667, A11004, A11011, A21050, SA5-10094). Samples were rinsed and washed prior to mounting. A similar protocol was used for pMad, fibrillarin, and HP1a staining, with the following exceptions. For pMad, samples were rinsed and washed in PBS with 0.1% Triton X-100 plus 0.2% Tween 20 (Sigma, P9416), incubated with 1:500 rabbit anti-pMad (Abcam #ab52903) for 6 to 8 hours at 4°C and with 1:1000 Alexa Fluor 568- conjugated secondary antibody for 45 minutes at room temperature instead. For fibrillarin and HP1a, we first incubated samples with 1:1000 mouse anti-fibrillarin (38F3) (Abcam #ab4566) or 1:500 mouse HP1a (DSHB #C1A9; deposited by L. L. Wallrath) followed by incubation with 1:400 Alexa Fluor 568-conjugated secondary antibodies, and samples were rinsed and washed before incubation with α-spectrin and lamin C primary antibodies followed by incubation with 1:400 Alexa Fluor 488-conjugated secondary antibodies. For EU staining, briefly, the ovaries were incubated for 1 hour at room temperature in 100 μM EU (Invitrogen, Click-iT® RNA, C10330) diluted in Grace’s insect medium, washed and fixed as described above. After fixation, samples were rinsed once and washed with 0.5% Triton X-100 in PBS for 15’ at room temperature before being subjected to the Click-iT reaction according to the manufacturer’s instructions (Invitrogen, Click-iT® RNA, C10330) for 30 minutes at room temperature. Ovaries were blocked and followed our standard incubation with antibodies prior to mounting. All samples were mounted in VectaShield mounting media with 4’,6-diamidino-2-phenylindole (DAPI) (Vector Labs, H-1200-10), and images were collected on an LSM 900 confocal system using the integrated Airyscan 2 detector and corresponding Airyscan SR mode (ZEISS Microscopy) and processed with the standard Airyscan Processing utilities of the ZEN Blue 3.5 software (ZEISS Microscopy) to achieve super-resolution readouts. For each experiment, the same microscope settings were used for image acquisition in all conditions.

### RNA fluorescence *in situ* hybridization

Ovaries were prepared for RNA fluorescence *in situ* hybridization using a previously described *R2* Stellaris probe set (123,146). Briefly, ovaries were dissected in Dulbecco’s PBS 1x (DPBS) (Gibco, 14190-136) on ice and incubated with fixation solution [4% formaldehyde methanol-free (FA) (Thermo Scientific) in DPBS 1x] for 30 minutes at room temperature, washed twice with DPBS at room temperature for 5 minutes each, and incubated with cold 70% Molecular Biology Grade ethanol (Thermo Scientific) in DPBS at 4°C overnight on a nutator. Samples were rinsed twice and washed once with wash buffer [20% v/v Stellaris RNA FISH Wash Buffer A (SMF- WA1-60, LGC, Biosearch Technologies) plus 70% v/v nuclease-free water (Thermo Scientific) plus 10% v/v deionized formamide (VWR Life Science, 0606)] before hybridization at 37°C overnight with *R2* Stellaris probes in custom 3’ amine oligos in plates (LGC, Biosearch Technologies, Petaluma, CA). Samples were washed three times with Wash Buffer A for 15 minutes each at room temperature and mounted in VectaShield with DAPI prior to confocal microscopy.

### Quantification of fluorescence images

We used three approaches to analyze raw microscopy images. For measuring the intensities and volumes of Fibrillarin and Udd and the number of *R2* and γ-H2Av foci, we used the “3D Objects Counter” plugin (147) in ImageJ (148), applying the same threshold to all images within the same experiment. For EU quantification, we used a single optical slice along the largest diameters of each follicle and manually outlined each nurse cell nucleus (based on DAPI) for intensity measurements of DAPI and EU channels, using ZEN Blue 3.5 software (ZEISS Microscopy). The final values were obtained after subtracting the background of each channel and calculating the ratio of EU to DAPI. Fold changes were determined based on the average for 25°C, which was set as 1. For HP1a quantification, we used all Z-stacks spanning GSC nuclei and manually outlined each nucleus (based on DAPI) using ImageJ. The intensities of DAPI and HP1a channels were measured, and final values and fold changes were calculated as described above. For pMad quantification, we used a single optical slice along the largest diameter of each GSC nucleus and manually outlined the nucleus (based on DAPI) to measure pMad average intensity per pixel using ZEN Blue 3.5 software (ZEISS Microscopy) and calculate fold changes relative to 25°C control.

## Acknowledgments

A.C.P.G. performed all sample preparations and experiments under D.D.-B.’s mentorship, and Z.G. did the TE expression re-analyses of the UW–Madison Biotechnology Core’s data under G.L.’s mentorship. A.C.P.G. and D.D.-B. wrote the manuscript. Antibodies against α-Spectrin, Lamin C, and HP1a were obtained from Developmental Studies Hybridoma Bank (DSHB), created by the NICHD and maintained at the University of Iowa, Iowa City, IA 52242. We are grateful to M. Buszczak for the Udd antibody; J. Nelson, for helping with the *R2* FISH protocol; M. Busche and J. Brunkard for the kind sharing of reagents, instrumentation, and advice for nanoplate dPCR experiments; and M. Stefely for help with design of diagrams shown in Fig. 1A- 1D. Finally, we thank R. Nunes and A. Williams for helpful edits and comments on the manuscript.

## Supporting Information captions

**S1 Fig. Warm and mild cold temperatures induce distinct splice variant-level transcriptomic responses.** (A) Principal component analysis graph showing intra-group clustering and intergroup separation and indicating splice-variant expression correlation among different temperature conditions. Elliptical shapes indicate 95% confidence interval. (B) Heatmap of splice variants significantly (*q*≤0.05) modulated by temperature. Expression level is scaled per splice variant in each row, with yellow and blue representing upregulation and downregulation, respectively.

**S2 Fig. Mild cold strongly downregulates genes involved in piRNA processing.** Heatmap showing piRNA processing genes that are significantly (*q*≤0.05) modulated by 29°C or 18°C. Yellow and blue represent upregulated and downregulated genes, respectively.

**S3 Fig. Mild cold alters the levels of chromatin marks in germ cells.** Single optical slices of ovarioles from females maintained at 25°C control (A, D, and G), 18°C (B, E, and H), or 29°C (C, F, and I) for 5 days. DAPI (magenta) labels nuclei. H3K9ac (A-C’), H3K9me3 (D-F’), and HP1a (G-I’) are labeled in white, with single channels shown in (A’, B’, and C’), (D’, E’, and F’), and (G’, H’, and I’), respectively. The differences shown in these images are representative of those consistently observed across ovarioles analyzed. Samples sizes per temperature: 120 ovarioles (A-C); 150 ovarioles (D-F); 300 ovarioles (G-I). Scale bar: 50 µm.

**S1 Table. Transcriptomics results S2 Table. Venn diagram analysis**

**S3 Table. Gene set enrichment analysis**

**S4 Table. Summary of gene set enrichment analysis**

**S5 Table. Analysis of differentially regulated transposable elements S1 File. Quality control results for transcriptomic analysis**

