## Supplementary figures and images for "Distinct transcriptional responses to mild cold versus warm temperatures in adult *Drosophila melanogaster* ovaries"

### Supplemental Figure 1

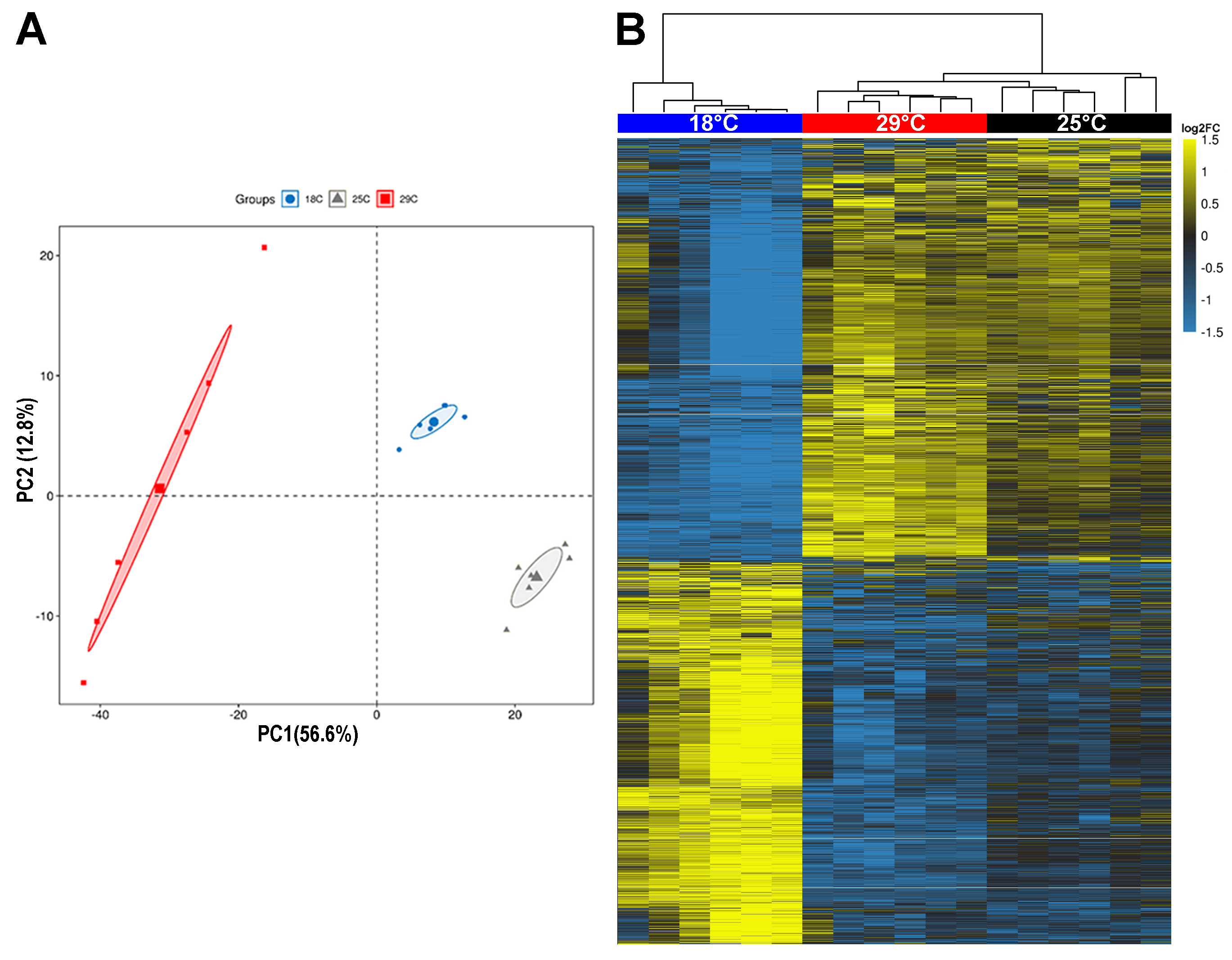

### Supplemental Figure 2

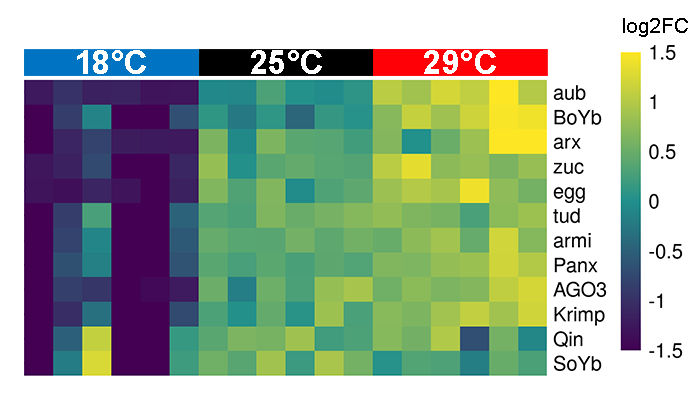

### Supplemental Figure 3

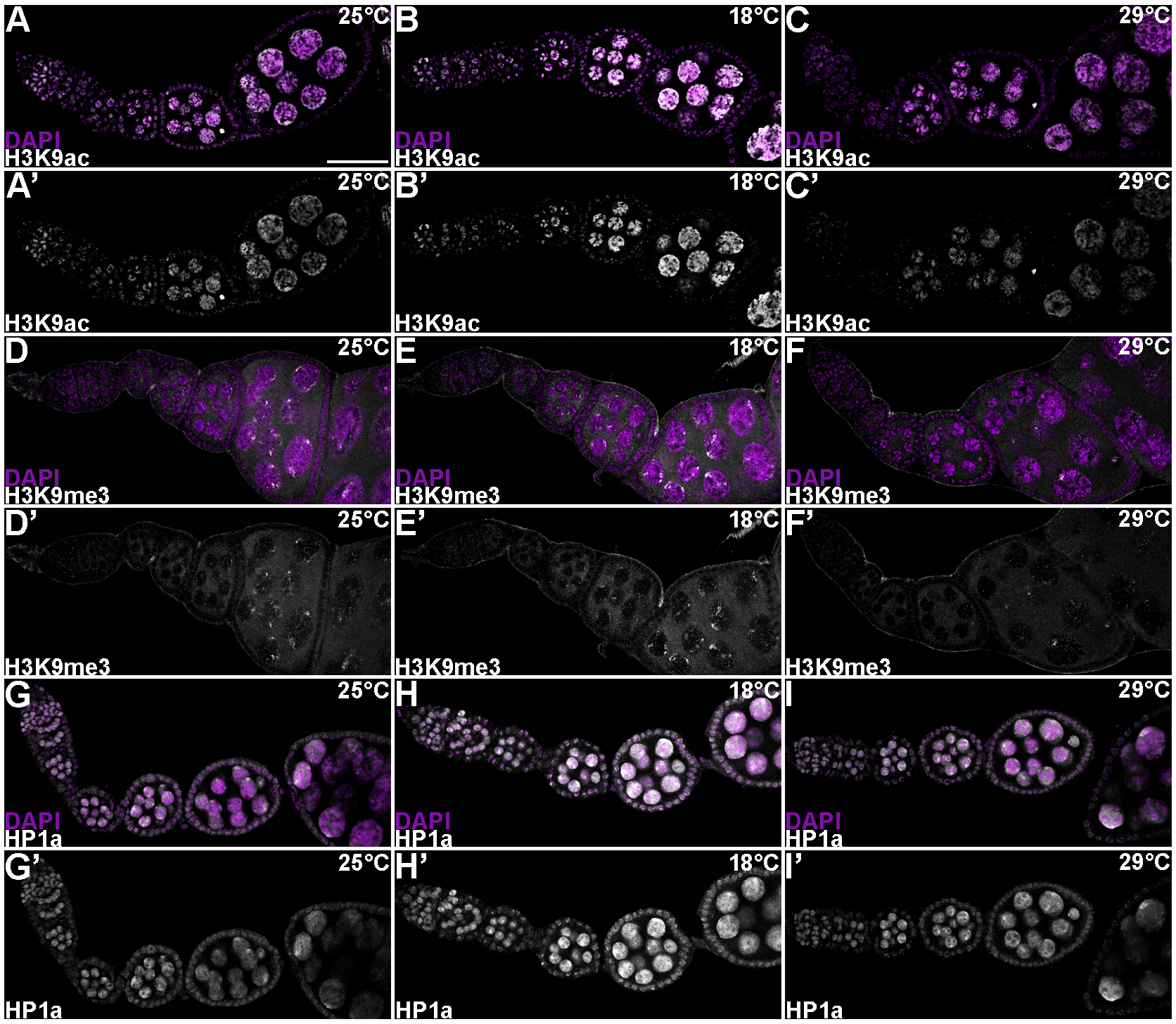
