## Supplemental Table 4 for "Distinct transcriptional responses to mild cold versus warm temperatures in adult *Drosophila melanogaster* ovaries"

**S4 Table. Summary of gene set enrichment analysis.**

| Temperature modulation | Top 10 GO terms "Biological Processes"[&] | General category[#] | Modulated genes in general category* |
|---|---|---|---|
| **Regulated in opposite directions at 29°C and 18°C** | GO:0071867 response to monoamine<br>GO:0071868 cellular response to monoamine stimulus<br>GO:0071869 response to catecholamine<br>GO:0071870 cellular response to catecholamine stimulus<br>GO:1903350 response to dopamine<br>GO:1903351 cellular response to dopamine<br>GO:0007212 G protein-coupled dopamine receptor signaling pathway<br>GO:0048499 synaptic vesicle membrane organization<br>GO:0008039 synaptic target recognition<br>GO:0007186 G protein-coupled receptor signaling pathway | Neuronal and GPCR signaling | ***Gbeta5***, ***CG42450***, ***sand***, ***Ten-a***, ***comt***, ***CG32373***, ***GABA-B-R3***, ***CG42450***, ***Proc-R***, *Galphas*, *Syt4*, *Ten-m*, *gek*, *Npc2a*, *CG9304*, *Camta*, *Lrpprc2*, *Octbeta2R*, *stops*, *fz*, *CG32638* |
| **Uniquely upregulated at 29°C** | GO:0007271 synaptic transmission, cholinergic<br>GO:0007268 chemical synaptic transmission<br>GO:0098916 anterograde trans-synaptic signaling<br>GO:0099537 trans-synaptic signaling<br>GO:0099536 synaptic signaling<br>GO:0034220 monoatomic ion transmembrane transport | Synaptic transmission and ion transport | ***nAChRbeta1***, ***Ace***, ***TkR99D***, *GABA-B-R2*, *Pka-R1*, *homer*, *Syx4*, *CG11155*, ***Porin2***, *CG6878*, *MME1* |
| | GO:0006334 nucleosome assembly<br>GO:0034728 nucleosome organization<br>GO:0065004 protein-DNA complex assembly<br>GO:0071824 protein-DNA complex organization | Nucleosome organization and genomic integrity | ***His1:CG33855***, ***His1:CG33858***, *His1:CG33834*, *His1:CG33855*, *spn-D* |
| **Uniquely downregulated at 29°C** | GO:0019367 fatty acid elongation, saturated fatty acid<br>GO:0019368 fatty acid elongation, unsaturated fatty acid<br>GO:0034625 fatty acid elongation, monounsaturated fatty acid<br>GO:0034626 fatty acid elongation, polyunsaturated fatty acid<br>GO:0030497 fatty acid elongation<br>GO:0042761 very long-chain fatty acid biosynthetic process<br>GO:0000038 very long-chain fatty acid metabolic process<br>GO:0006633 fatty acid biosynthetic process<br>GO:0072330 monocarboxylic acid biosynthetic process | Fatty acid biosynthesis | ***eloF***, ***CG16904***, *sit*, *Sc2*, ***CG15531***, ***FASN3***, *FASN1*, *CG42233* |

| | | | |
|---|---|---|---|
| | GO:0030148 sphingolipid biosynthetic process | | |
| **Uniquely upregulated at 18°C** | GO:1903341 regulation of meiotic DNA double-strand break formation<br>GO:1903343 positive regulation of meiotic DNA double-strand break formation<br>GO:0042138 meiotic DNA double-strand break formation | Meiotic DNA double-strand break formation | ***vilya***, ***narya*** |
| | GO:0060048 cardiac muscle contraction<br>GO:0071689 muscle thin filament assembly<br>GO:0006941 striated muscle contraction<br>GO:0060361 flight<br>GO:0006936 muscle contraction<br>GO:0034315 regulation of Arp2/3 complex-mediated actin nucleation | Muscle function and actin cytoskeleton | ***up***, ***KCNQ***, ***Mhc***, ***Mlc2***, ***Tm2***, ***Mlc1***, ***nwk***, ***PICK1***, *tn*, *Fkbp12*, *Chd64*, *wupA*, *SCAR*, *Arfip*, *GMF* |
| | GO:2000378 negative regulation of reactive oxygen species metabolic process | Reactive oxygen species metabolism | ***Vdup1***, ***Tace*** |
| **Uniquely downregulated at 18°C** | GO:0042066 perineurial glial growth<br>GO:0042065 glial cell growth | Glial growth | ***Nf1***, ***poe*** |
| | GO:0051382 kinetochore assembly<br>GO:0034508 centromere complex assembly<br>GO:0051383 kinetochore organization<br>GO:0007080 mitotic metaphase chromosome alignment | Kinetochore organization | ***Cenp-C, Spc105R, nudE***, *cal1, Cam, asp* |
| | GO:0016339 calcium-dependent cell-cell adhesion via plasma membrane cell adhesion molecules<br>GO:0044331 cell-cell adhesion mediated by cadherin | Cell-cell adhesion and synaptic transmission | ***kug***, ***Cals***, ***CG1909***, mys, Cad87A, CadN, *beta-Spec*, *chico*, *cv-c*, *CaMKII* |
| | GO:0050806 positive regulation of synaptic transmission<br>GO:1905954 positive regulation of lipid localization | Regulation of lipid localization | ***Apoltp***, ***Lsd-1***, *LpR2*, *Pi3K21B*, *InR*, *Acsl*, *bmm*, *Itpr*, *Pi3K92E* |

[&]GO terms in grey font are not significantly enriched among all modulated genes.
[#]Fits most genes from combined GO terms.
*Genes in bold are, at least, 2-fold regulated.
