## Supplemental file 1 for "Distinct transcriptional responses to mild cold versus warm temperatures in adult *Drosophila melanogaster* ovaries"

A. Absorbance

### RNA 10/10/2023 11:04 AM : Samples

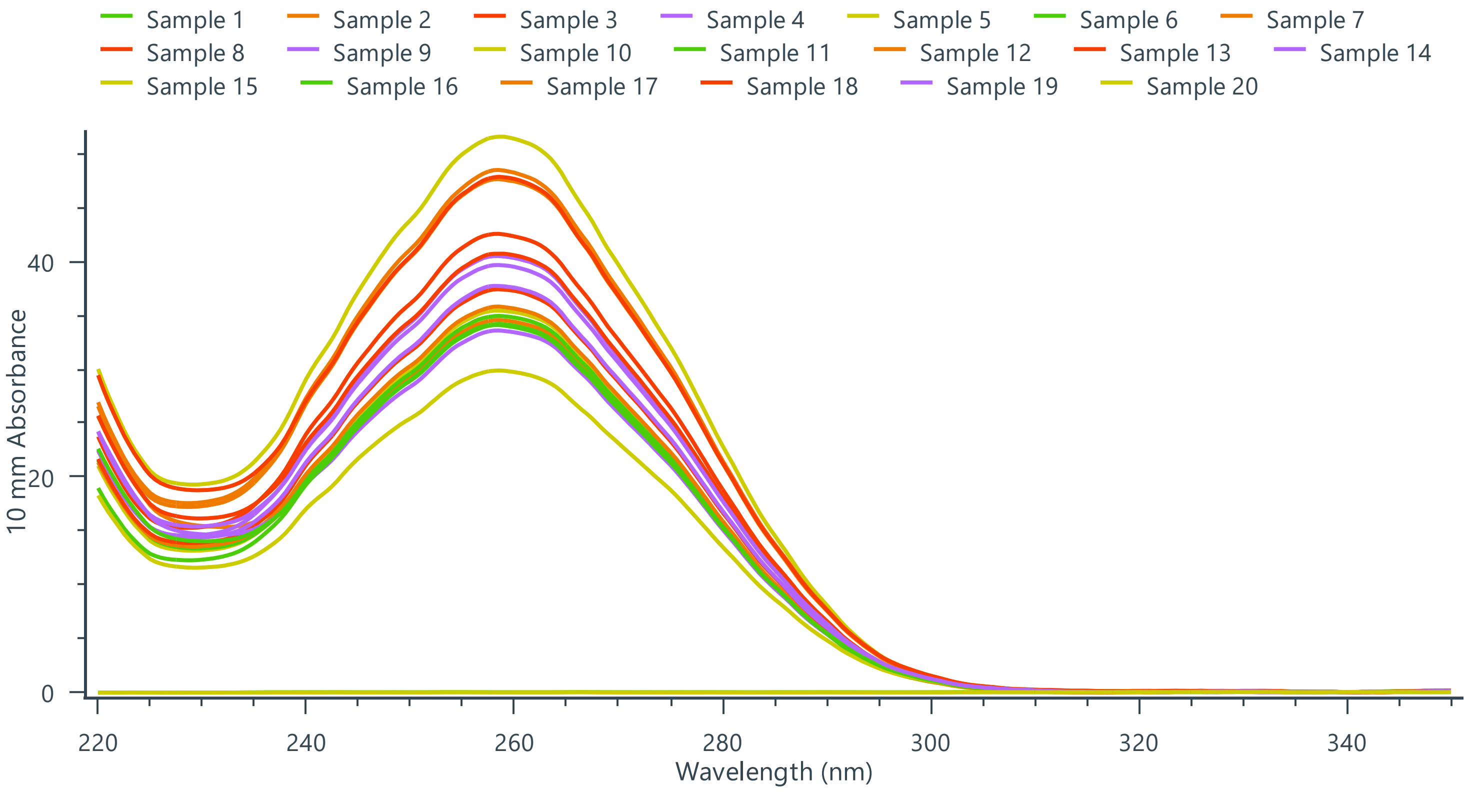

B. Nanodrop quantification

### Results

| Name | Date | Conc(ng/µL) | A260/A280 | A260/A230 | A260 | A280 |
|---|---|---|---|---|---|---|
| Sample 1 | 10/10/2023 11:04:24 AM | -0.6 | 1.24 | 0.37 | -0.02 | -0.01 |
| Sample 2 | 10/10/2023 11:05:06 AM | 1376.3 | 2.23 | 2.23 | 34.41 | 15.41 |
| Sample 3 | 10/10/2023 11:05:39 AM | 1693.6 | 2.25 | 2.76 | 42.34 | 18.84 |
| Sample 4 | 10/10/2023 11:06:06 AM | 1613.1 | 2.25 | 2.76 | 40.33 | 17.96 |
| Sample 5 | 10/10/2023 11:06:37 AM | 1412.2 | 2.25 | 2.68 | 35.31 | 15.72 |
| Sample 6 | 10/10/2023 11:07:15 AM | 1392.0 | 2.24 | 2.61 | 34.80 | 15.55 |
| Sample 7 | 10/10/2023 11:07:44 AM | 1425.5 | 2.24 | 2.63 | 35.64 | 15.92 |
| Sample 8 | 10/10/2023 11:08:13 AM | 1491.3 | 2.23 | 2.68 | 37.28 | 16.69 |
| Sample 9 | 10/10/2023 11:08:40 AM | 1337.3 | 2.21 | 2.28 | 33.43 | 15.12 |
| Sample 10 | 10/10/2023 11:09:09 AM | 2056.3 | 2.25 | 2.66 | 51.41 | 22.89 |
| Sample 11 | 10/10/2023 11:09:42 AM | 1356.3 | 2.22 | 2.75 | 33.91 | 15.27 |
| Sample 12 | 10/10/2023 11:10:11 AM | 1898.1 | 2.23 | 2.75 | 47.45 | 21.27 |
| Sample 13 | 10/10/2023 11:10:40 AM | 1623.4 | 2.23 | 2.52 | 40.59 | 18.20 |
| Sample 14 | 10/10/2023 11:11:05 AM | 1503.3 | 2.23 | 2.62 | 37.58 | 16.85 |
| Sample 15 | 10/10/2023 11:11:31 AM | 1189.3 | 2.20 | 2.57 | 29.73 | 13.50 |
| Sample 16 | 10/10/2023 11:11:58 AM | 1361.7 | 2.21 | 2.43 | 34.04 | 15.39 |
| Sample 17 | 10/10/2023 11:12:25 AM | 1930.5 | 2.24 | 2.74 | 48.26 | 21.52 |
| Sample 18 | 10/10/2023 11:12:54 AM | 1907.3 | 2.24 | 2.54 | 47.68 | 21.32 |
| Sample 19 | 10/10/2023 11:13:21 AM | 1580.4 | 2.22 | 2.56 | 39.51 | 17.76 |

| Name | Date | Conc(ng/µL) | A260/A280 | A260/A230 | A260 | A280 |
|---|---|---|---|---|---|---|
| Sample 20 | 10/10/2023 11:13:58 AM | -0.8 | 0.76 | 0.44 | -0.02 | -0.03 |

#### C. Electroforesis



**Filename: 2023-10-12 - 13.53.44.D1000**

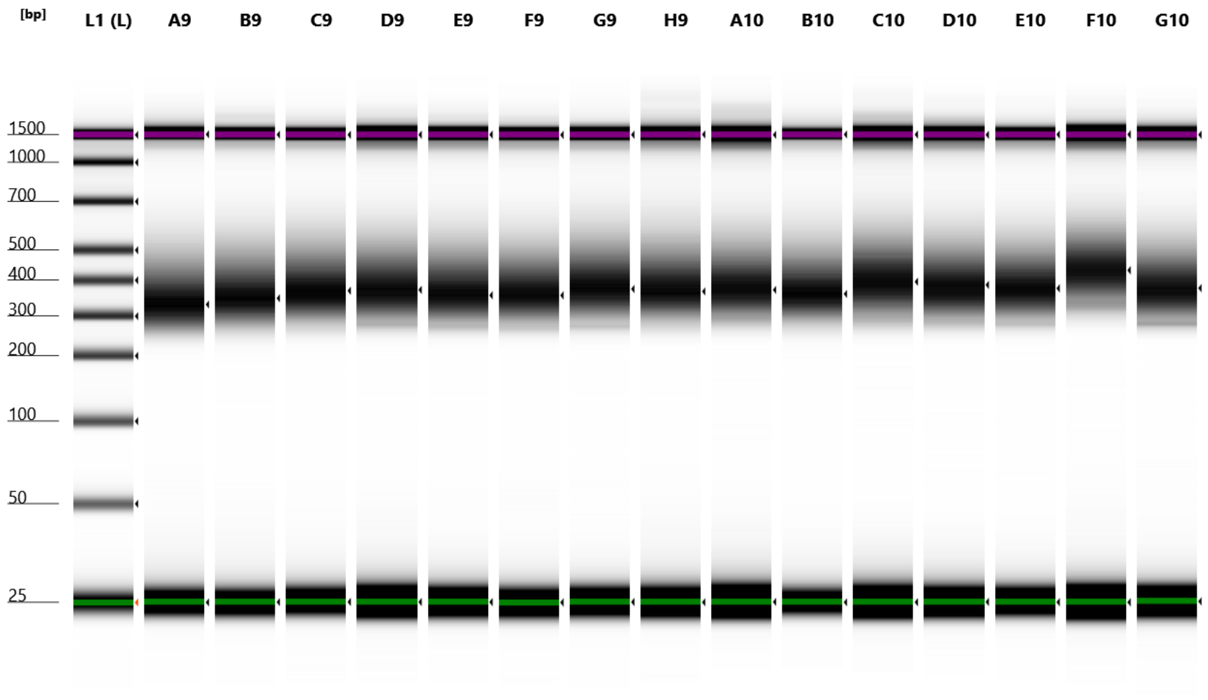


Default image (Contrast 50%), Image is Scaled to Highest Sample Peak

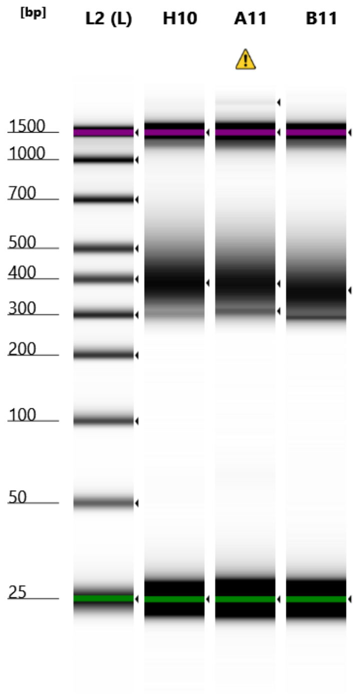


Default image (Contrast 50%), Image is Scaled to Highest Sample Peak


© Agilent Technologies, Inc. 2023

### D. RNA screentape data



#### Peak: L1: Ladder

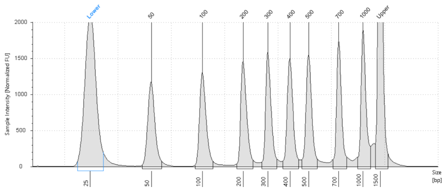

**Location** L1
**Concentration** 20.7
**Description** Ladder
**Observations** Ladder

| Size [bp] | Calibrated Conc. [ng/µl] | Assigned Conc. [ng/µl] | Peak Molarity [nmol/l] | % Integrated Area | Peak Comment | Observations |
|---|---|---|---|---|---|---|
| 25 | 6.90 | - | 425 | - | | Lower Marker |
| 50 | 2.50 | - | 76.8 | 12.06 | | |
| 100 | 2.53 | - | 38.9 | 12.23 | | |
| 200 | 2.57 | - | 19.8 | 12.41 | | |
| 300 | 2.58 | - | 13.2 | 12.48 | | |
| 400 | 2.60 | - | 9.99 | 12.55 | | |
| 500 | 2.66 | - | 8.18 | 12.85 | | |
| 700 | 2.52 | - | 5.55 | 12.20 | | |
| 1000 | 2.74 | - | 4.21 | 13.23 | | |
| 1500 | 6.50 | 6.50 | 6.67 | - | | Upper Marker |

#### Region: A9: 1-18C

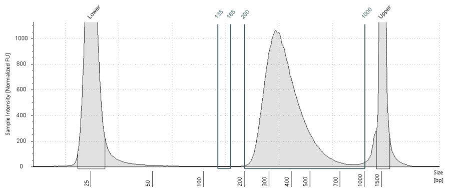

**Location** A9
**Concentration** 8.36
**Description** 1-18C
**Observations**

| From [bp] | To [bp] | Average Size [bp] | Conc. [ng/µl] | Region Molarity [nmol/l] | % of Total | Region Comment | Color |
|---|---|---|---|---|---|---|---|
| 135 | 165 | 151 | 0.000842 | 0.0237 | 0.01 | | |
| 200 | 1000 | 378 | 9.85 | 42.3 | 89.04 | | |

#### Region: B9: 2-18C

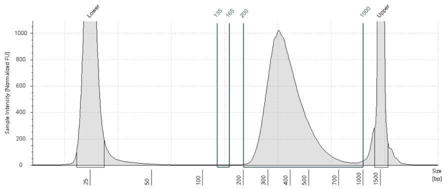

**Location** B9
**Concentration** 8.33
**Description** 2-18C
**Observations**

| From [bp] | To [bp] | Average Size [bp] | Conc. [ng/µl] | Region Molarity [nmol/l] | % of Total | Region Comment | Color |
|---|---|---|---|---|---|---|---|
| 135 | 165 | 152 | 0.000165 | 0.0111 | 0.00 | | |
| 200 | 1000 | 396 | 9.84 | 40.4 | 89.62 | | |

#### Region: C9: 3-18C

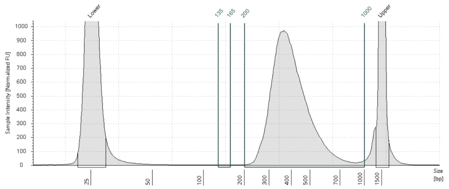

**Location** C9
**Concentration** 8.15
**Description** 3-18C
**Observations**

| From [bp] | To [bp] | Average Size [bp] | Conc. [ng/µl] | Region Molarity [nmol/l] | % of Total | Region Comment | Color |
|---|---|---|---|---|---|---|---|
| 135 | 165 | 152 | 0.000172 | 0.00957 | 0.00 | | |
| 200 | 1000 | 411 | 9.73 | 38.4 | 90.15 | | |


© Agilent Technologies, Inc. 2023

#### Region: D9: 4-18C

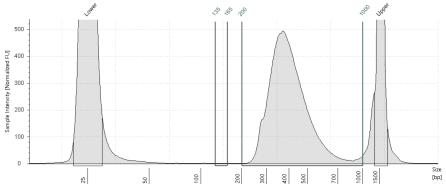

**Location** D9
**Concentration** 4.61
**Description** 4-18C
**Observations**

| From [bp] | To [bp] | Average Size [bp] | Conc. [ng/µl] | Region Molarity [nmol/l] | % of Total | Region Comment | Color |
|---|---|---|---|---|---|---|---|
| 135 | 165 | 153 | 0.000503 | 0.0205 | 0.01 | | |
| 200 | 1000 | 413 | 5.27 | 20.7 | 84.43 | | |

#### Region: E9: 5-18C

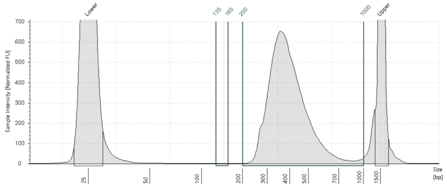

**Location** E9
**Concentration** 5.63
**Description** 5-18C
**Observations**

| From [bp] | To [bp] | Average Size [bp] | Conc. [ng/µl] | Region Molarity [nmol/l] | % of Total | Region Comment | Color |
|---|---|---|---|---|---|---|---|
| 135 | 165 | 149 | 0.000666 | 0.0242 | 0.01 | | |
| 200 | 1000 | 405 | 6.71 | 26.9 | 88.21 | | |

#### Region: F9: 6-18C

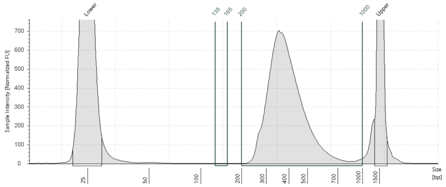

**Location** F9
**Concentration** 6.30
**Description** 6-18C
**Observations**

| From [bp] | To [bp] | Average Size [bp] | Conc. [ng/µl] | Region Molarity [nmol/l] | % of Total | Region Comment | Color |
|---|---|---|---|---|---|---|---|
| 135 | 165 | 150 | 0.00146 | 0.0251 | 0.02 | | |
| 200 | 1000 | 403 | 7.23 | 29.1 | 88.91 | | |

#### Region: G9: 7-25C

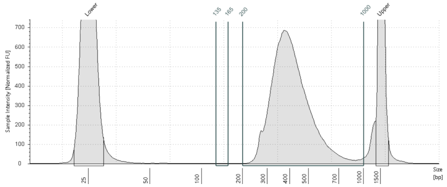

**Location** G9
**Concentration** 6.33
**Description** 7-25C
**Observations**

| From [bp] | To [bp] | Average Size [bp] | Conc. [ng/µl] | Region Molarity [nmol/l] | % of Total | Region Comment | Color |
|---|---|---|---|---|---|---|---|
| 135 | 165 | 149 | 0.00270 | 0.0382 | 0.03 | | |
| 200 | 1000 | 421 | 7.68 | 29.6 | 90.52 | | |



© Agilent Technologies, Inc. 2023

**Region: H9: 8-25C**

| | |
|---|---|
| **Location** | H9 |
| **Concentration** | 5.41 |
| **Description** | 8-25C |
| **Observations** | |

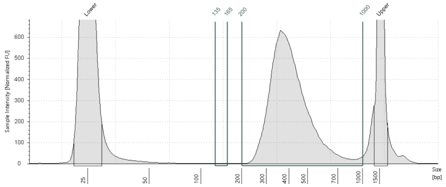

| From [bp] | To [bp] | Average Size [bp] | Conc. [ng/µl] | Region Molarity [nmol/l] | % of Total | Region Comment | Color |
|---|---|---|---|---|---|---|---|
| 135 | 165 | 151 | 0.00251 | 0.0391 | 0.03 | | |
| 200 | 1000 | 420 | 6.72 | 26.0 | 84.49 | | |

**Region: A10: 9-25C**

| | |
|---|---|
| **Location** | A10 |
| **Concentration** | 3.88 |
| **Description** | 9-25C |
| **Observations** | |

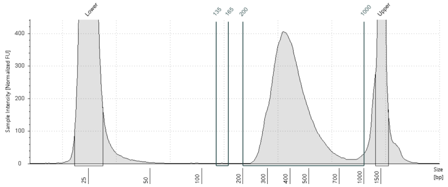

| From [bp] | To [bp] | Average Size [bp] | Conc. [ng/µl] | Region Molarity [nmol/l] | % of Total | Region Comment | Color |
|---|---|---|---|---|---|---|---|
| 135 | 165 | 151 | 0.00154 | 0.0315 | 0.03 | | |
| 200 | 1000 | 421 | 4.55 | 17.6 | 80.55 | | |

**Region: B10: 10-25C**

| | |
|---|---|
| **Location** | B10 |
| **Concentration** | 9.25 |
| **Description** | 10-25C |
| **Observations** | |

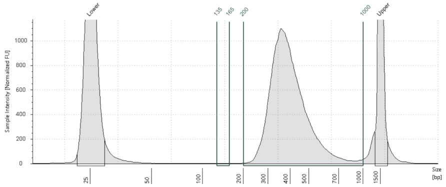

| From [bp] | To [bp] | Average Size [bp] | Conc. [ng/µl] | Region Molarity [nmol/l] | % of Total | Region Comment | Color |
|---|---|---|---|---|---|---|---|
| 135 | 165 | 151 | 0.00191 | 0.0324 | 0.02 | | |
| 200 | 1000 | 407 | 11.2 | 44.8 | 90.25 | | |

**Region: C10: 11-25C**

| | |
|---|---|
| **Location** | C10 |
| **Concentration** | 4.45 |
| **Description** | 11-25C |
| **Observations** | |

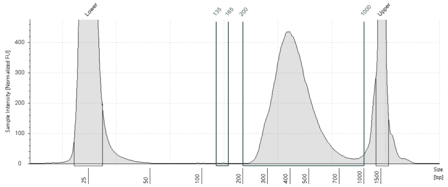

| From [bp] | To [bp] | Average Size [bp] | Conc. [ng/µl] | Region Molarity [nmol/l] | % of Total | Region Comment | Color |
|---|---|---|---|---|---|---|---|
| 135 | 165 | 150 | 0.00211 | 0.0343 | 0.03 | | |
| 200 | 1000 | 433 | 5.27 | 19.8 | 82.51 | | |


© Agilent Technologies, Inc. 2023

#### Region: D10: 12-25C

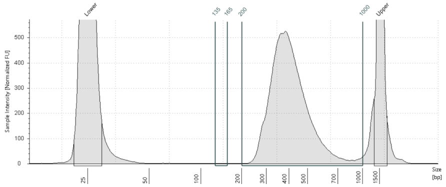

**Location** D10
**Concentration** 2.91
**Description** 12-25C
**Observations**

| From [bp] | To [bp] | Average Size [bp] | Conc. [ng/µl] | Region Molarity [nmol/l] | % of Total | Region Comment | Color |
|---|---|---|---|---|---|---|---|
| 135 | 165 | 151 | 0.000633 | 0.0160 | 0.01 | | |
| 200 | 1000 | 417 | 5.86 | 22.8 | 85.47 | | |

#### Region: E10: 13-29C

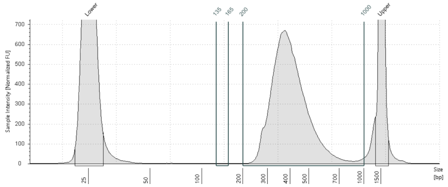

**Location** E10
**Concentration** 6.13
**Description** 13-29C
**Observations**

| From [bp] | To [bp] | Average Size [bp] | Conc. [ng/µl] | Region Molarity [nmol/l] | % of Total | Region Comment | Color |
|---|---|---|---|---|---|---|---|
| 135 | 165 | 153 | 0.00141 | 0.0240 | 0.02 | | |
| 200 | 1000 | 412 | 7.12 | 27.9 | 89.32 | | |

#### Region: F10: 14-29C

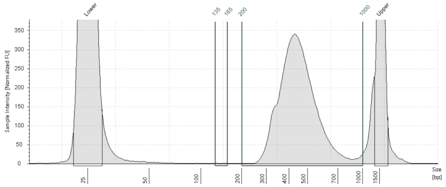

**Location** F10
**Concentration** 3.36
**Description** 14-29C
**Observations**

| From [bp] | To [bp] | Average Size [bp] | Conc. [ng/µl] | Region Molarity [nmol/l] | % of Total | Region Comment | Color |
|---|---|---|---|---|---|---|---|
| 135 | 165 | 151 | 0.00171 | 0.0275 | 0.04 | | |
| 200 | 1000 | 461 | 3.83 | 13.4 | 82.42 | | |

#### Region: G10: 15-29C

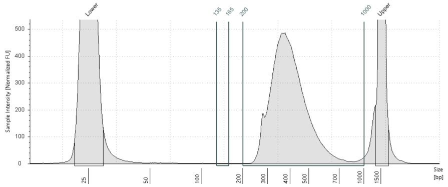

**Location** G10
**Concentration** 4.38
**Description** 15-29C
**Observations**

| From [bp] | To [bp] | Average Size [bp] | Conc. [ng/µl] | Region Molarity [nmol/l] | % of Total | Region Comment | Color |
|---|---|---|---|---|---|---|---|
| 135 | 165 | 153 | 0.000405 | 0.0136 | 0.01 | | |
| 200 | 1000 | 408 | 5.27 | 20.9 | 87.08 | | |


© Agilent Technologies, Inc. 2023

#### Peak: L2: Ladder

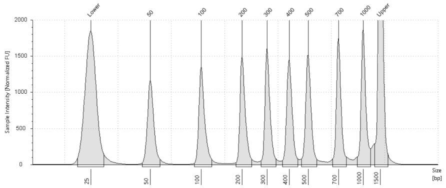

| | |
|---|---|
| **Location** | L2 |
| **Concentration** | 19.8 |
| **Description** | Ladder |
| **Observations** | Ladder |

| Size [bp] | Calibrated Conc. [ng/µl] | Assigned Conc. [ng/µl] | Peak Molarity [nmol/l] | % Integrated Area | Peak Comment | Observations |
|---|---|---|---|---|---|---|
| 25 | 6.39 | - | 393 | - | | Lower Marker |
| 50 | 2.29 | - | 70.4 | 11.57 | | |
| 100 | 2.41 | - | 37.1 | 12.20 | | |
| 200 | 2.45 | - | 18.8 | 12.38 | | |
| 300 | 2.49 | - | 12.8 | 12.61 | | |
| 400 | 2.46 | - | 9.44 | 12.41 | | |
| 500 | 2.55 | - | 7.86 | 12.91 | | |
| 700 | 2.47 | - | 5.44 | 12.51 | | |
| 1000 | 2.65 | - | 4.08 | 13.41 | | |
| 1500 | 6.50 | 6.50 | 6.67 | - | | Upper Marker |

#### Region: H10: 16-29C

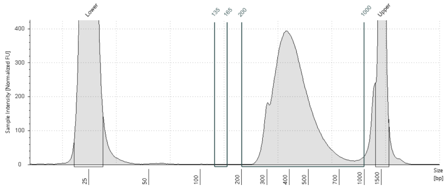

| | |
|---|---|
| **Location** | H10 |
| **Concentration** | 3.47 |
| **Description** | 16-29C |
| **Observations** | |

| From [bp] | To [bp] | Average Size [bp] | Conc. [ng/µl] | Region Molarity [nmol/l] | % of Total | Region Comment | Color |
|---|---|---|---|---|---|---|---|
| 135 | 165 | 155 | 0.000383 | 0.0107 | 0.01 | | |
| 200 | 1000 | 431 | 4.35 | 16.4 | 84.87 | | |

#### Region: A11: 17-29C

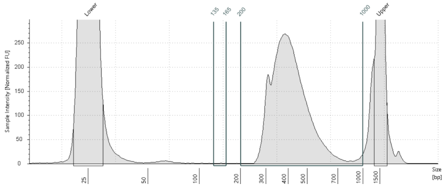

| | |
|---|---|
| **Location** | A11 |
| **Concentration** | 2.71 |
| **Description** | 17-29C |
| **Alert** | ⚠ |
| **Observations** | Peak out of Sizing Range |

| From [bp] | To [bp] | Average Size [bp] | Conc. [ng/µl] | Region Molarity [nmol/l] | % of Total | Region Comment | Color |
|---|---|---|---|---|---|---|---|
| 135 | 165 | 154 | 0.00125 | 0.0216 | 0.03 | | |
| 200 | 1000 | 422 | 3.06 | 11.7 | 79.26 | | |

#### Region: B11: 18-29C

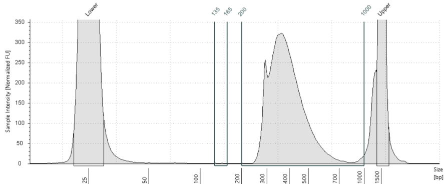

| | |
|---|---|
| **Location** | B11 |
| **Concentration** | 2.75 |
| **Description** | 18-29C |
| **Observations** | |

| From [bp] | To [bp] | Average Size [bp] | Conc. [ng/µl] | Region Molarity [nmol/l] | % of Total | Region Comment | Color |
|---|---|---|---|---|---|---|---|
| 135 | 165 | 149 | 0.00121 | 0.0218 | 0.03 | | |
| 200 | 1000 | 399 | 3.58 | 14.5 | 81.72 | | |


© Agilent Technologies, Inc. 2023

© Agilent Technologies, Inc. 2023

GENES

E. Boxplot of normalized counts

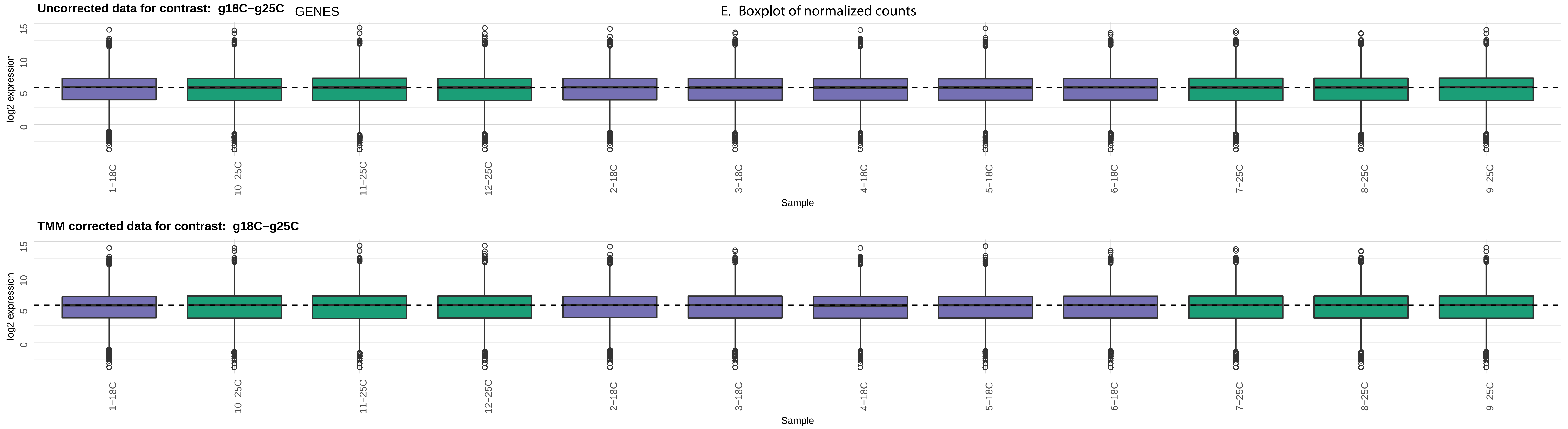

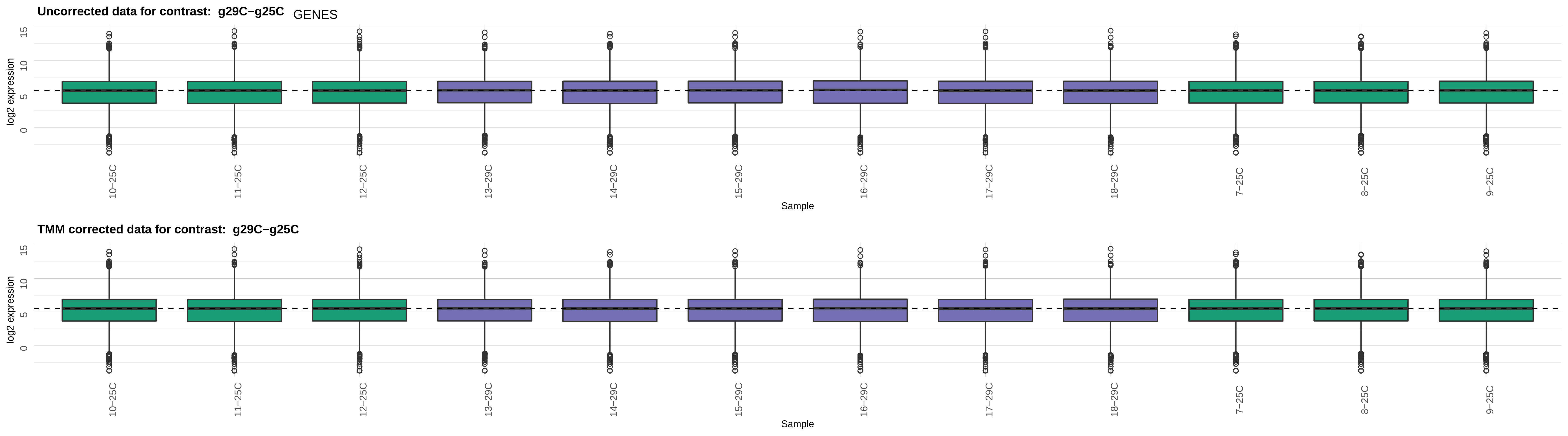
Uncorrected data for contrast: g29C−g25C
GENES
log2 expression
0
5
10
15
10−25C
11−25C
12−25C
13−29C
14−29C
15−29C
16−29C
17−29C
18−29C
7−25C
8−25C
9−25C
Sample
TMM corrected data for contrast: g29C−g25C
log2 expression
0
5
10
15
10−25C
11−25C
12−25C
13−29C
14−29C
15−29C
16−29C
17−29C
18−29C
7−25C
8−25C
9−25C
Sample

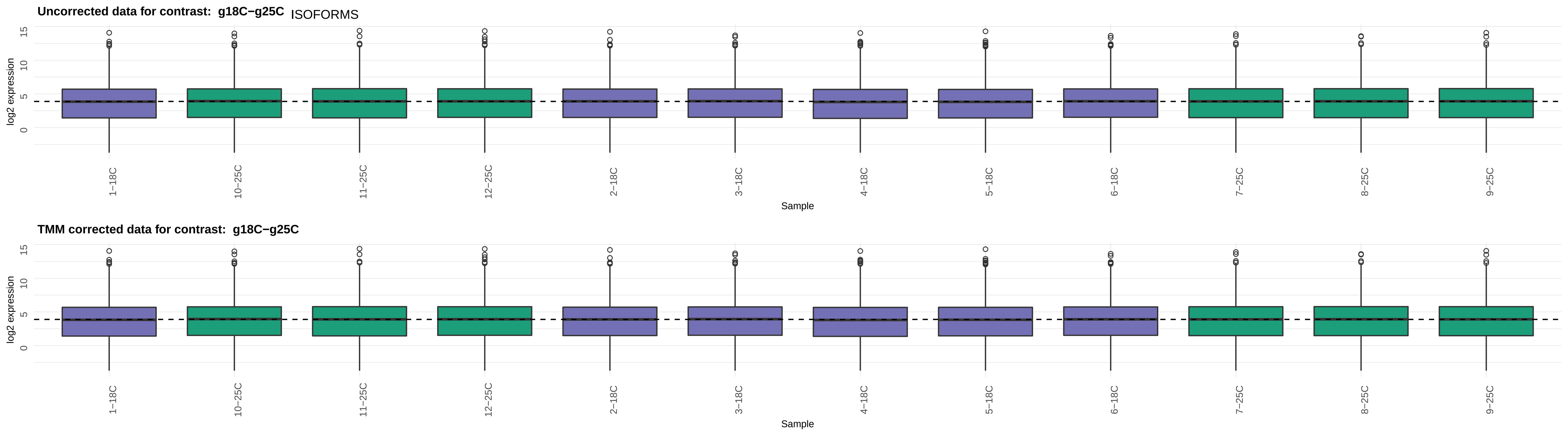
Uncorrected data for contrast: g18C−g25C
ISOFORMS
log2 expression
15
10
5
0
1−18C
10−25C
11−25C
12−25C
2−18C
3−18C
4−18C
5−18C
6−18C
7−25C
8−25C
9−25C
Sample
TMM corrected data for contrast: g18C−g25C
log2 expression
15
10
5
0
1−18C
10−25C
11−25C
12−25C
2−18C
3−18C
4−18C
5−18C
6−18C
7−25C
8−25C
9−25C
Sample

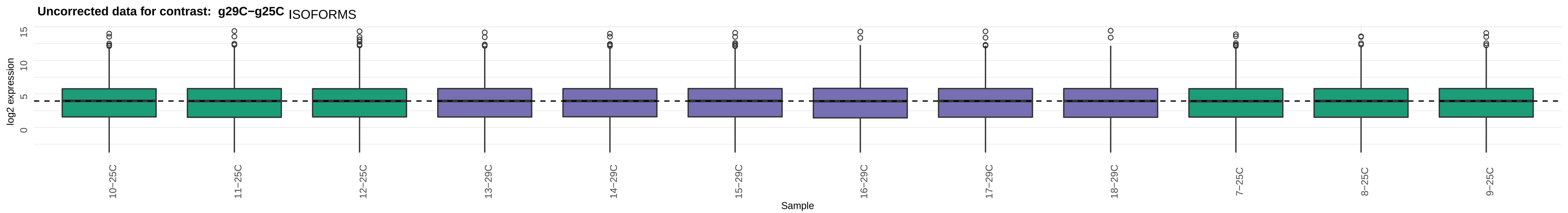

Uncorrected data for contrast: g29C−g25C
ISOFORMS
log2 expression
0
5
10
15
10−25C
11−25C
12−25C
13−29C
14−29C
15−29C
16−29C
17−29C
18−29C
7−25C
8−25C
9−25C
Sample


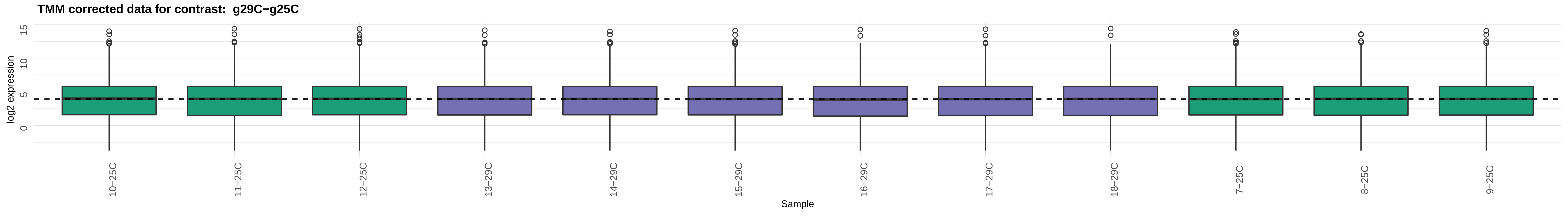

TMM corrected data for contrast: g29C−g25C
log2 expression
0
5
10
15
10−25C
11−25C
12−25C
13−29C
14−29C
15−29C
16−29C
17−29C
18−29C
7−25C
8−25C
9−25C
Sample
